# The polar flagellar transcriptional regulatory network in *Vibrio campbellii* deviates from canonical *Vibrio* species

**DOI:** 10.1101/2021.05.10.443453

**Authors:** Blake D. Petersen, Michael S. Liu, Ram Podicheti, Albert Ying-Po Yang, Chelsea Simpson, Chris Hemmerich, Douglas B. Rusch, Julia C. van Kessel

**Affiliations:** Department of Biology, Indiana University, Bloomington, IN 47405; Center for Genomics and Bioinformatics, Indiana University, Bloomington, IN 47405; Graduate Institute of Oncology, College of Medicine, National Taiwan University, Taipei 100, Taiwan

**Keywords:** motility, flagellar gene cluster, flagellar gene regulation, *Vibrio*, *Vibrio campbellii*

## Abstract

*Vibrio campbellii* is a Gram-negative bacterium that is free-living and ubiquitous in marine environments, and it is a pathogen of fish and shellfish. Swimming motility via a single polar flagellum is a critical virulence factor in *V. campbellii* pathogenesis, and disruption of the flagellar motor significantly decreases host mortality. To examine *V. campbellii* flagellar gene regulation, we identified homologs of flagellar and chemotaxis genes conserved in other members of the *Vibrionaceae* and determined the transcriptional profile of these loci using differential RNA-seq. We systematically deleted all 63 predicted flagellar and chemotaxis genes in *V. campbellii* and examined their effects on motility and flagellum production. We specifically focused on the core flagellar regulators of the flagellar regulatory hierarchy established in other *Vibrios*: RpoN (σ^54^), FlrA, FlrC, and FliA. Our results show that *V. campbellii* transcription of flagellar and chemotaxis genes is governed by a multi-tiered regulatory hierarchy similar to other motile *Vibrio* species but with two critical differences: the σ^54^-dependent regulator FlrA is dispensable for motility, and Class II gene expression is independent of σ^54^ regulation. Our genetic and phenotypic dissection of the *V. campbellii* flagellar regulatory network highlights the differences that have evolved in flagellar regulation across the *Vibrionaceae*.

## Introduction

Swimming motility is an important behavior that is common among aquatic bacteria. By using a rotating molecular nanomachine called a flagellum, bacteria can propel themselves through fluid environments as a resource-scavenging strategy (1, 2). For many bacteria swimming motility is also critical for host interactions. *Vibrio* species are a highly motile group of marine bacteria that closely associate with a diverse range of marine hosts (3–6). Some vibrios form symbiotic pairings, such the mutualism between *Vibrio fischeri* and the Hawaiian bobtail squid, *Euprymna scolopes* (3, 7). Other vibrios are pathogenic and can infect humans (*e*.*g*., *Vibrio cholerae, Vibrio parahaemolyticus*) and/or are pathogens of fish and shellfish (*e*.*g*., *Vibrio harveyi, Vibrio campbellii*) (8–11). Flagellar-mediated motility plays a critical role for each of these interactions between *Vibrio* species and their hosts; both colonization and lethality of these strains in their respective hosts decreases when motility is inhibited (4, 12, 13). For this reason, vibrios have served as key models for study of flagellar regulation and its role in colonization.

Of the various *Vibrio* species, *V. cholerae* has been the most studied *Vibrio* species regarding flagellar regulation and has thus served as the predominant model for polar flagellar regulation in other vibrios (14–16). In *V. cholerae*, flagellar regulation is organized stepwise into a four-tiered transcriptional hierarchy, whereby expression of downstream flagellar components is activated after expression of upstream components (16, 17). Under this model, the sole Class I gene encodes the σ^54^-dependent transcriptional activator FlrA. Once FlrA is expressed, Class II gene expression is activated, which includes several structural components of the flagellum (flagellar export machinery, early basal body genes, ATPase), some chemotaxis genes, and the *flrBC* and *fliA* regulatory genes whose products regulate expression of Class III and IV, respectively (16, 18, 19). Class III is activated by FlrC, another σ^54^-dependent transcriptional regulator that belongs to a two-component system along with its cognate histidine kinase FlrB (19, 20). Class III genes include intermediate structural components, such as the distal and proximal rod, hook proteins, and the essential flagellin, *flaA* (16). Once the flagellum has reached a certain length after Class III expression, the anti-sigma factor FlgM is exported from the cell, leading to activation of FliA (the alternative sigma factor σ^28^) that activates Class IV genes (21). These include late structural components such as the flagellar stator, four alternative flagellins, the T ring, as well as other chemotaxis proteins, including the methyl-accepting chemoreceptors. The synthesis of a functional flagellum in *V. cholerae* depends on this stepwise regulatory hierarchy, and deletion of any of the four regulators disrupts the subsequent steps and results in complete loss of motility (19).

The regulatory network driving flagellar motility has also been studied in other *Vibrio* species, including *V. fischeri, V. parahaemolyticus*, and *Vibrio vulnificus* (22). The polar flagellar genes are highly conserved between *Vibrio* species and are organized into similar genetic clusters in each genome (16, 22, 23). Despite this, flagellar phenotypes differ among each species. While most *Vibrios* produce a monotrichous polar sheathed flagellum, some species such as *V. fischeri* exhibit peritrichous polar sheathed flagella (24, 25). *V. parahaemolyticus* and *Vibrio alginolyticus* encode a secondary set of flagellar genes that allow them to produce peritrichous lateral unsheathed flagella in addition to their polar flagellum, enabling them to undergo swarming motility over surfaces (26, 27). These differences in motile phenotypes suggest that there is much diversity in the number, type, and regulation of flagella among *Vibrio* species.

*V. campbellii* is a marine bacterium belonging to the Harveyi clade that shares similar free-living and pathogenic life cycles with other members of the *Vibrionaceae* (28, 29). *V. campbellii* has served as an important organism for the study of pathogenesis because it infects many different organisms critical to aquaculture, including shrimp, oysters, and fish (8–10). *V. campbellii* uses a monotrichous sheathed polar flagellum to propel itself. As with other vibrios, swimming motility plays an important role in pathogenesis for *V. campbellii*, and inhibition of motility significantly decreases host mortality during infection (13). However, while *V. campbellii* encodes flagellar and chemotaxis genes conserved by other *Vibrio species*, the expression and function of these genes have not been characterized. In addition, the *V. campbellii* genome encodes genes with homology to *V. parahaemolyticus* lateral flagellar genes for swarming motility. However, only a few strains appear to express these genes with little phenotypic or genetic characterization (30, 31). Thus, there are several gaps in our understanding of *V. campbellii* flagellar-based motility.

Recently, we established the wild isolate *V. campbellii* DS40M4 as a new model strain for *V. campbellii* research because it is capable of natural transformation, thus facilitating powerful genetic manipulations such as MuGENT (32–34). The genetic tractability of DS40M4 enabled us to use it as a model system to study flagellar motility in *V. campbellii*. We systematically deleted all 63 predicted polar flagellar and chemotaxis genes in *V. campbellii* and characterized the swimming phenotypes of each mutant. We used RNA-seq to identify the genes controlled by canonical flagellar regulators in *V. campbellii*. With these data, we propose a model for the transcriptional regulation program that governs expression of polar flagellar genes in *V. campbellii*.

## Results

### *Identification of the flagellar gene operons in* V. campbellii

Using reciprocal BLASTp analyses based on known flagellar homologs characterized in other Vibrios, we identified 63 distinct flagellar genes organized across five regions within Chromosome I of *V. campbellii* DS40M4 (Fig. 1A) (22). In our previous work, we conducted differential RNA-seq (dRNA-seq) to identify transcription start sites across the entire DS40M4 genome (33). We used the dRNA-seq data to identify transcriptional start sites in these five loci, allowing us to predict corresponding operons for each flagellar gene (Fig. 1A). Examples of the dRNA-seq transcript data are shown in Figure 1B for the *fliEFGHIJ, pomAB*, and *flhAFG* genes (Fig. 1B). Thus, Figure 1 presents a diagram showing the predicted organization of the core polar flagellar and chemotaxis genes in *V. campbellii* and respective transcriptional start sites for each. (Fig. 1).

**Figure 1:**
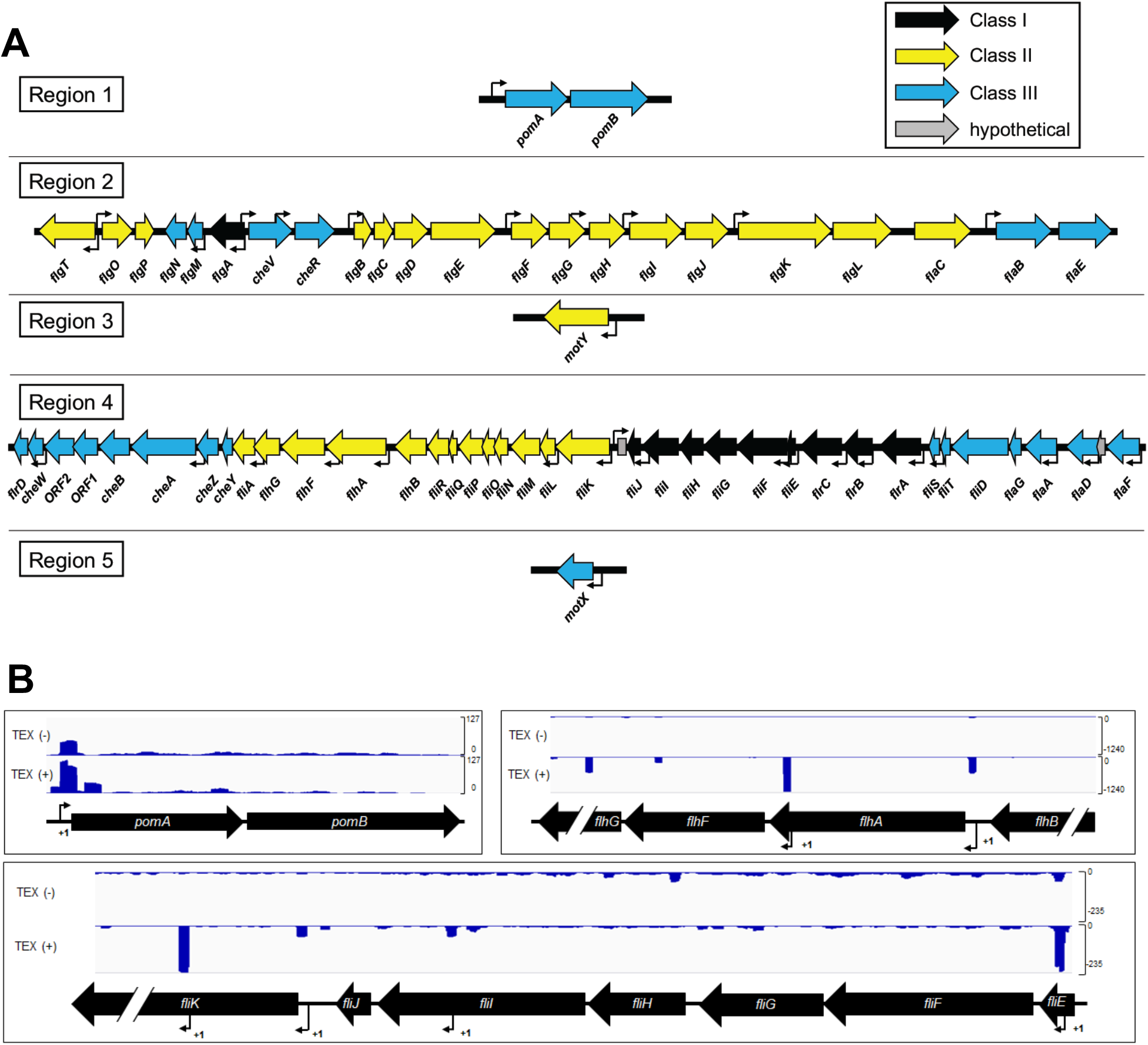
Organization of the polar flagellar gene system in *V. campbellii* DS40M4 A) Diagram showing the five regions of polar flagellar genes in Chromosome I in *V. campbellii*. Transcriptional start sites (TSS) are mapped to each region (black arrows) based on dRNA-seq results. Flagellar genes are color coded based on their classification in the transcriptional hierarchy via our RNA-seq data. B) Data shown are cDNA reads from dRNA-seq data mapped to three genetic loci in *V. campbellii* DS40M4 for samples untreated (TEX (-)) or treated (TEX (+)) with terminator 5’-phosphate-dependent endonuclease. Small black arrows labeled with “ +1” indicate predicted TSS for flagellar genes.

We next used comparative genomics to investigate the conservation of each of the known polar flagellar genes between *V. campbellii* and other known polar flagellated *Vibrio* species (Fig. 2). We used a minimum amino acid percent identity requirement of 40% (the lowest possible value for clustering allowed by cd-hit) in this analysis to increase the number of functional homologs identified in our results. For the flagellins that are highly similar to each other, we also independently examined their similarity to flagellins in other *Vibrio* species using BLASTp. Our genomic data suggest that the genome of *V. campbellii* encodes homologs that match the complete set of known polar flagellar genes established in *Vibrio* species thus far (Fig. 2). These genes are highly conserved in the *V. campbellii* clade. The flagellar gene system in *V. campbellii* shares amino acid identity most closely with *V. parahaemolyticus*, whereas most polar flagellar genes in *V. campbellii* are more divergent compared to other members of the *Vibrionaceae*, such as *V. cholerae* or *V. fischeri*.

**Figure 2:**
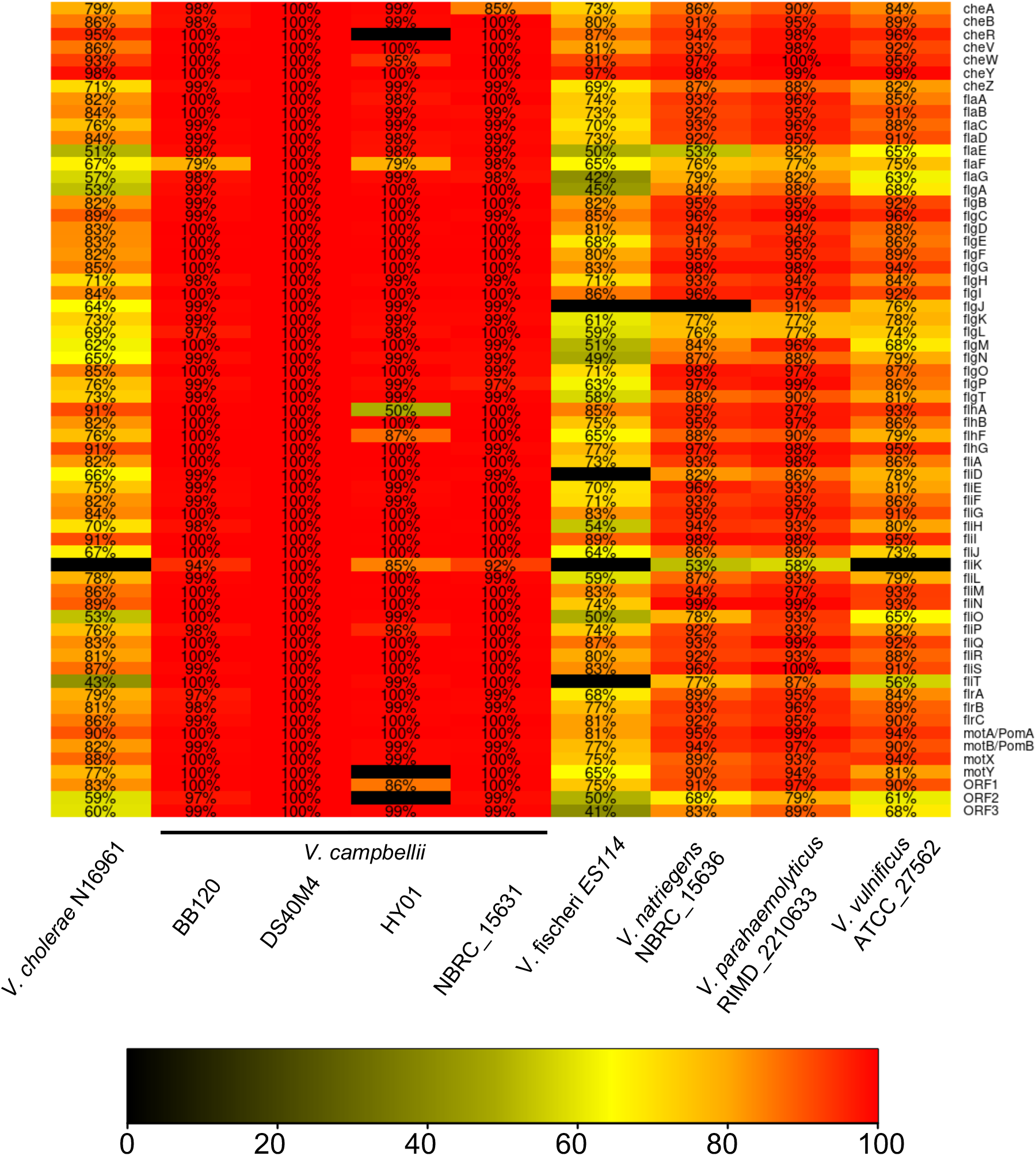
Polar flagellar gene homology across *Vibrionaceae*. Comparative genomics showing amino acid identity of known *V. cholerae* polar flagellar genes. This chart indicates the amino acid identity shared with *V. campbellii* DS40M4 proteins as indicated by the color scale.

### *Deletion and characterization of all known flagellar/chemotaxis genes in* V. campbellii

Few studies have characterized the complete set of flagellar genes in a *Vibrio* species, likely owing to the time and labor associated with genetic mutation of dozens of genes (22, 23). Recently we showed that genetic manipulation of *V. campbellii* DS40M4 is highly efficient due to inducible natural transformation via overexpression of the master competence regulator Tfox (32, 33). To automate primer design for constructing deletion mutants in *V. campbellii*, we programmed an algorithm to generate a list of primers for deletion of every annotated open reading frame in the DS40M4 genome. Candidate primers were designed so that overlapping genes were not disrupted and followed a specific set of criteria for validity, including preference for shorter length (25-30 nucleotides), no homopolymers of five or more bases, a melting temperature of 58-63°C, and the 3’ end of each primer ends in a cytosine or guanine. We used this algorithm to design primers to construct unmarked deletions of each predicted flagellar or chemotaxis gene in *V. campbellii*.

Altogether, we constructed 64 mutants, including each of the 63 flagellar and chemotaxis genes spread across the five flagellar regions in Chromosome I of DS40M4, as well as *rpoN* that encodes σ^54^. We classified mutants into groups based on their capacity to swim via soft agar swim plates and the presence or absence of flagella observed via microscopy using the fluorescence stain NanoOrange, a reagent which is virtually nonfluorescent in aqueous solution that fluoresces brightly when it binds hydrophobic regions of proteins and lipids, allowing visualization of cell bodies and flagella (35, 36). Mutants that deviated from wild-type were classified into three phenotypic classes: (1) nonmotile (Mot^-^) mutants are inhibited for motility on swim plates but produce a detectable flagellum; (2) aflagellate (Fla^-^) mutants are inhibited for motility on swim plates and produce no detectable flagellum; (3) semimotile (Mot^±^) mutants demonstrate reduced motility (smaller swim halos) on swim plates (Table 1). Of these 64 mutants, 34 are still capable of producing a swim halo (ranging from significantly reduced to wild-type levels), while 30 are completely inhibited for swimming motility on swim plates (Fig. 3A). The reduced motility phenotypes in some of these mutants may be explained by a chemotaxis defect such as in *che* genes (i.e. *cheABRWVYZ*), which were not studied further within the scope of this work. Overall, we identified a total of 7 Mot^-^ mutants, 25 Fla^-^ mutants, and 20 Mot^±^ mutants, with 12 mutants that have phenotypes indistinguishable from wild-type.

**Table 1.**
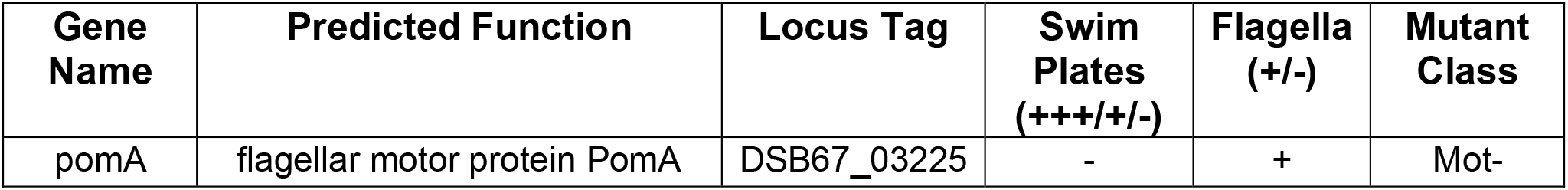
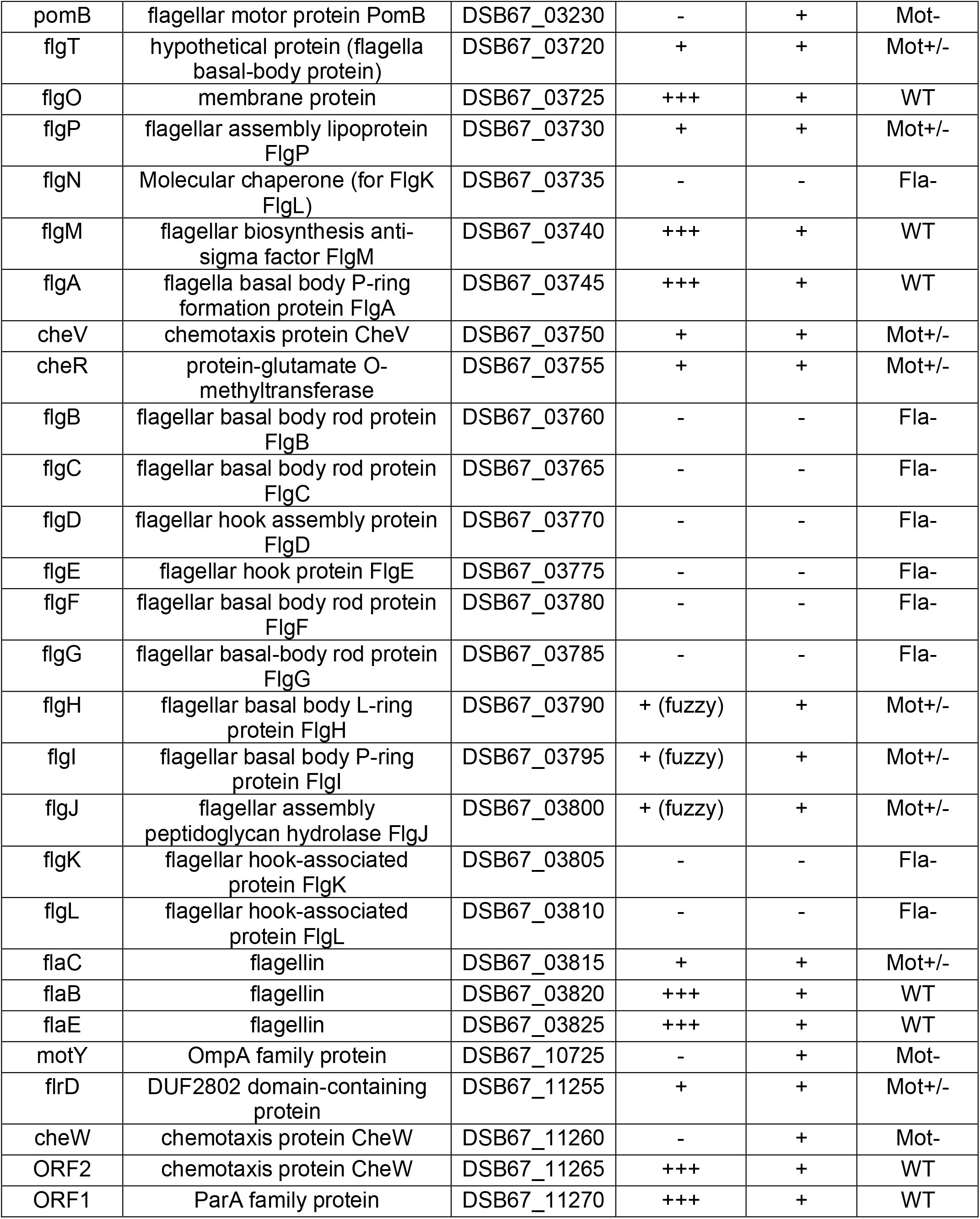
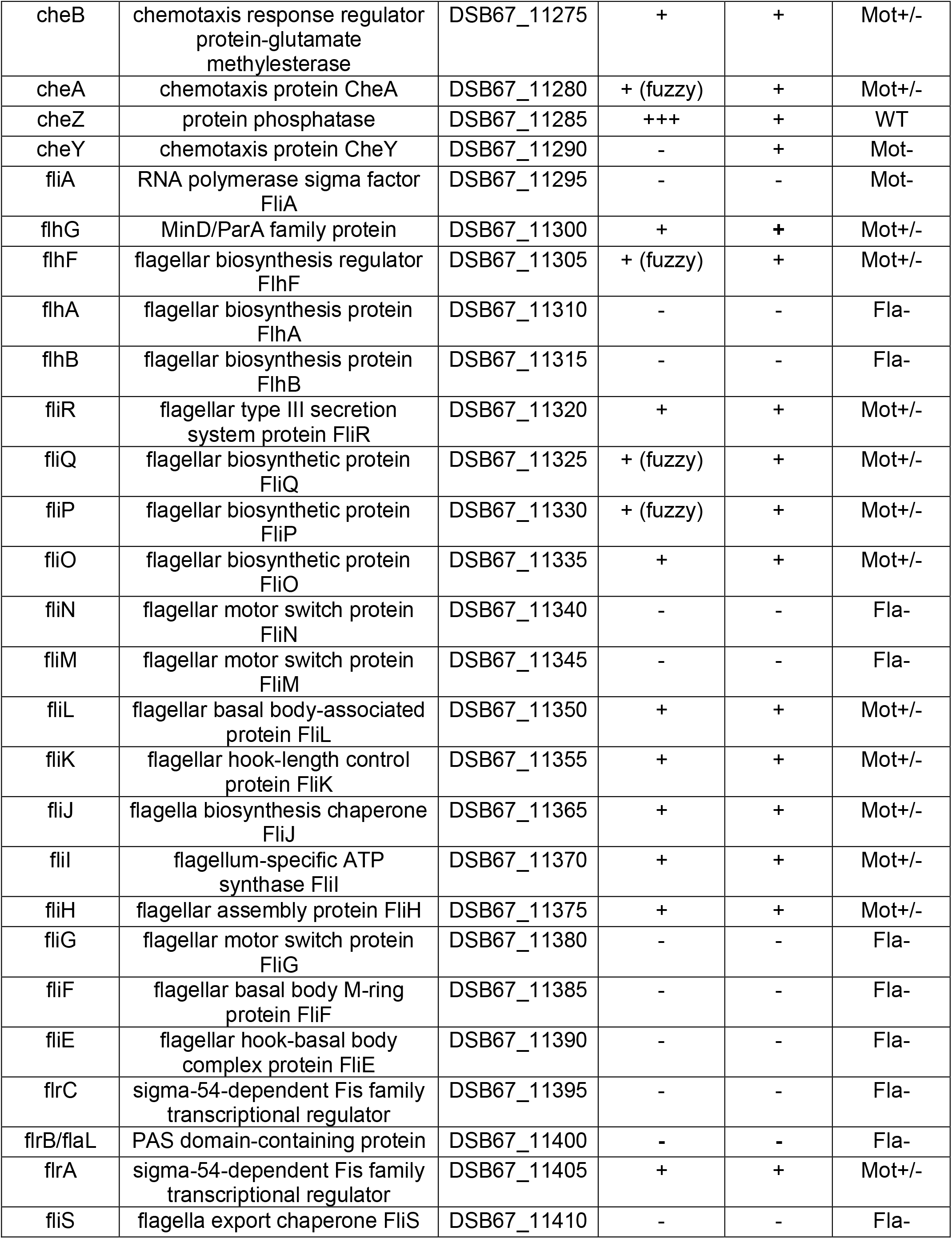
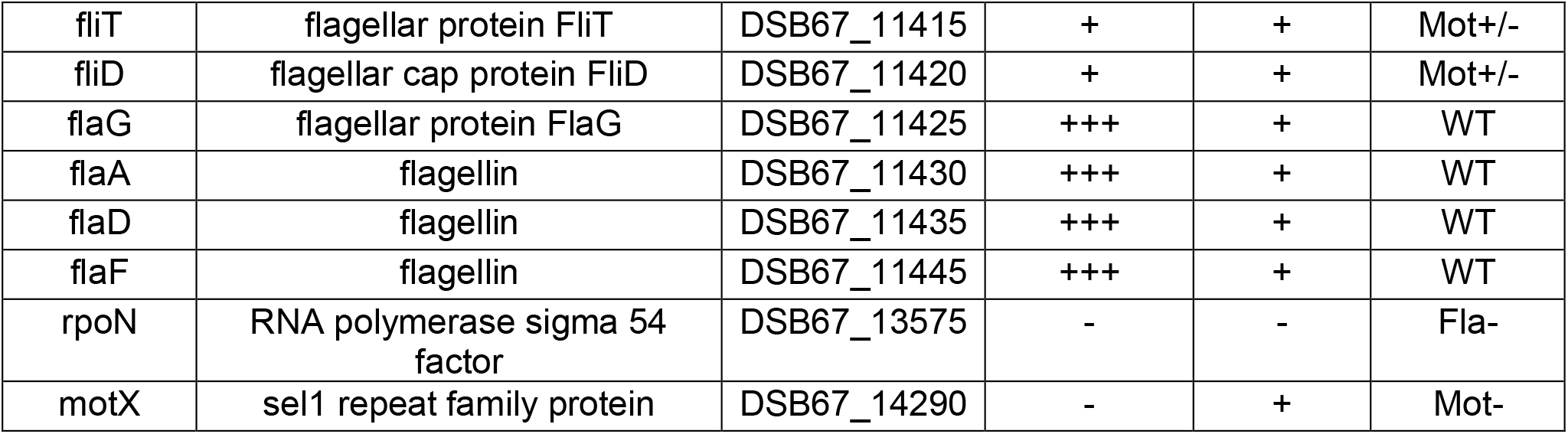
Phenotypic classification of *V. campbellii* DS40M4 strains containing deletions of predicted flagellar or chemotaxis genes.

**Figure 3.**
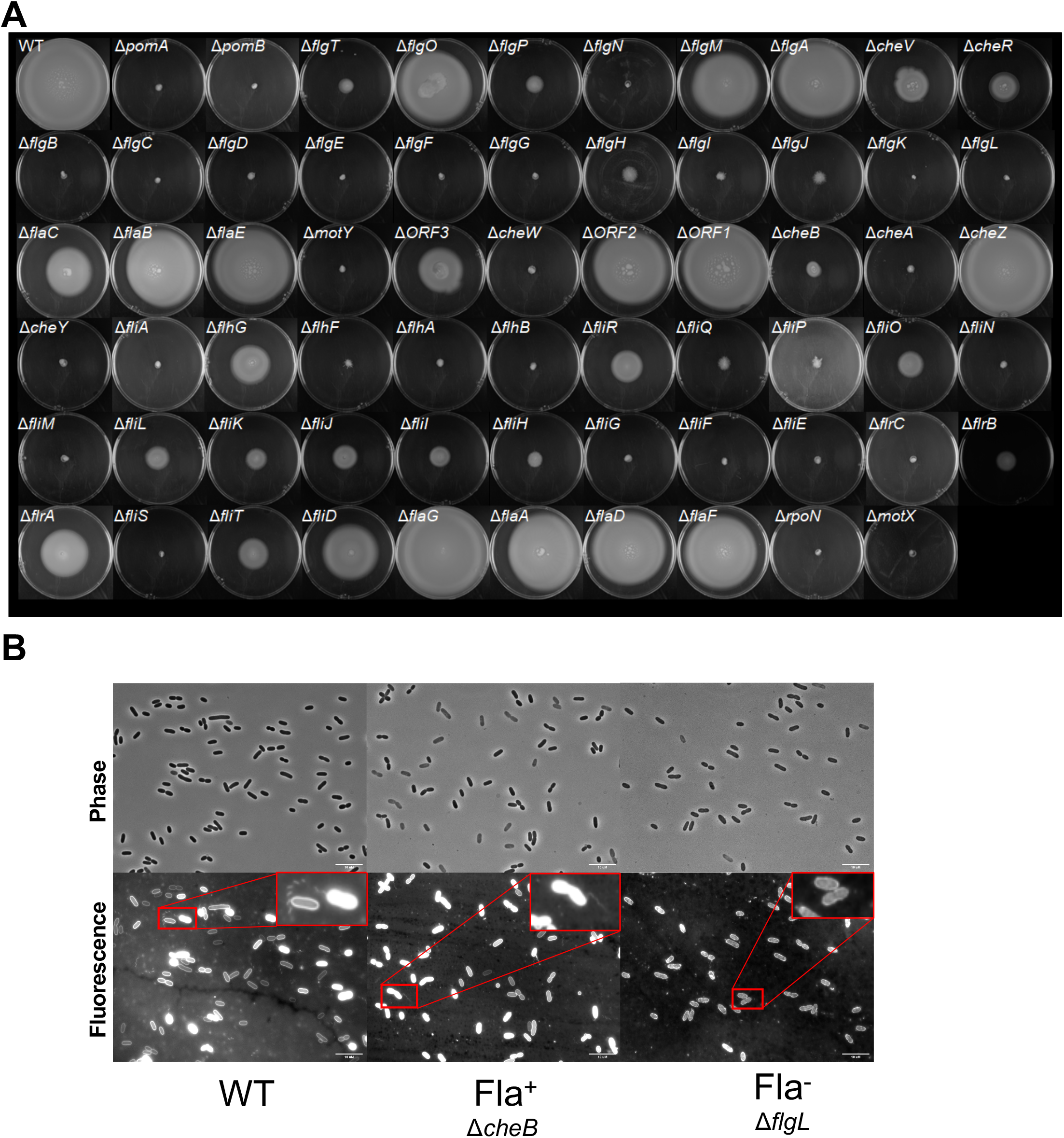
Swim phenotypes of DS40M4 polar flagellar mutants. A) Soft agar (0.3%) swim plates showing swim halo phenotypes for wild-type and mutant strains of DS40M4. B) Fluorescence microscopy images of wild-type, D*cheB* (Fla+), and D*flgL* (Fla-) strains using the fluorescent protein stain NanoOrange to observe the presence of flagella.

Among the 7 Mot^-^ mutants from our pool are the four motor/stator proteins *pomAB, motX*, and *motY*, which are crucial for flagellar rotation but not assembly, as well as *fliA* (σ^28^; Class IV regulator) *and cheYW* (chemotaxis) (CITE). The 25 Fla^-^ mutants included *flgN* (hook chaperone), *flgBCF* (proximal rod), *flgDE* (hook), *flgG* (distal rod), *flgH* (L ring), *flgI* (P ring), *flgJ* (rod cap), *flgKL* (hook), *flhBA* (exporter apparatus), *fliEF* (MS ring), *fliGMN* (C ring), *fliPQ* (exporter apparatus), *fliS* (flagellin chaperone), *flrBC* (Class III regulators), and σ^54^. Among the mutants in our pool of 20 Mot^±^ mutants, we identified *flgT* and *flgP* (H ring), *cheVR* (chemotaxis), *flaC* (flagellin), *flrD* (unknown), *cheAB* (chemotaxis), *flhFG* (flagellar number/localization), *fliHIJ* (ATPase), *fliK* (hook length), *fliL* (torque generator), *fliOR* (export apparatus), *flrA* (Class II regulator), *fliT* (chaperone), and *fliD* (filament cap). Finally, we found little to no change in motility in *flgM* (anti-σ^28^), *flgO* (H ring), *flgA, flaABDEF* (flagellins), *cheZ* (chemotaxis), and ORF1/ORF2 (unknown/chemotaxis) mutants compared to the wild-type strain.

Most mutants within our pool of 64 were sorted into phenotypic classes that agree with previous observations in other vibrios or flagellate bacterial species. However, some discrepancies were found compared to these findings. Notable examples include mutants in the flagellar export apparatus (*fliOR*) and the filament cap (*fliD* and *fliT*), both of which are required for motility in other bacteria such as *Salmonella enterica* but resulted in Mot^±^ phenotypes for our mutants (37–40). Although soft agar plates are often used to assess swimming motility, the interpretation of these can be misleading because the observed phenotypes are the result of a combination of factors including but not limited to flagellar synthesis, chemotactic sensing, growth rates, biofilm formation, and sensing of metabolites. Thus, in addition to using swim plates we used phase-contrast microscopy to directly observe swimming in all semimotile mutants. We distinguished semimotile mutants by creating a fourth phenotypic class that we designated nonswimmers (Swm^-^), which produce swim halos on soft agar plates but show no detectable swimming cells when observed by live cell microscopy. While we observed swimming in many of the Mot^±^ mutants, we were surprised to identify several mutants that were semimotile on soft agar plates appear nonmotile by microscopy, including *flgT* (H ring), *flrD* (regulatory protein), *flhF* (flagellar placement), *fliO* (export apparatus), and *fliHIJ* (ATPase). These results corroborate previous observations of flagellar mutants in other bacteria, in which deletion of these genes results in nonmotile and/or aflagellate cells (37, 41–43). This suggests an important difference between the behaviors observed on swim plates versus microscope that may be explained by defects in flagellar assembly, structure, or function.

### Effects of core flagellar transcriptional regulators on swimming motility and flagellum synthesis

In vibrios with a defined polar flagellar gene system, including *V. cholerae, V. fischeri, V. vulnificus*, and *V. parahaemolyticus*, previous studies have shown that the flagellum is regulated by a four-tiered transcriptional hierarchy that includes σ^54^, FlrA (or the homolog FlaK), FlrC (or the homolog FlaM), and FliA (σ^28^). Deletion of any one of these transcriptional regulators results in complete loss of motility in most vibrios, with the exception of *V. parahaemolyticus* that also produces a compensating regulator of lateral flagella, LafK (18, 19, 44–46). To determine if the *V. campbellii* flagellar genes are regulated in a similar transcriptional hierarchy, we examined the four predicted regulators: σ^54^, FlrA (annotated as FlaK), FlrC (annotated as FlaM), and FliA. While the annotation of *V. campbellii* DS40M4 shares the same naming scheme for FlaKLM with *V. parahaemolyticus*, we refer to them as FlrABC, respectively, henceforth for consistency with the naming in other vibrios and due to the meaning behind the prefix Flr (<u>fl</u>agellar regulatory protein) (44). We constructed isogenic strains with a single deletion of each of the flagellar transcriptional regulator genes in DS40M4 and tested for the presence or absence of motility in each mutant on soft agar plates (Fig. 3A). Similar to other vibrios, deletion of *rpoN* (that encodes *σ*^*54*^), *flrC*, and *fliA* results in the complete abrogation of motility on swim plates (Fig 4A).

**Figure 4.**
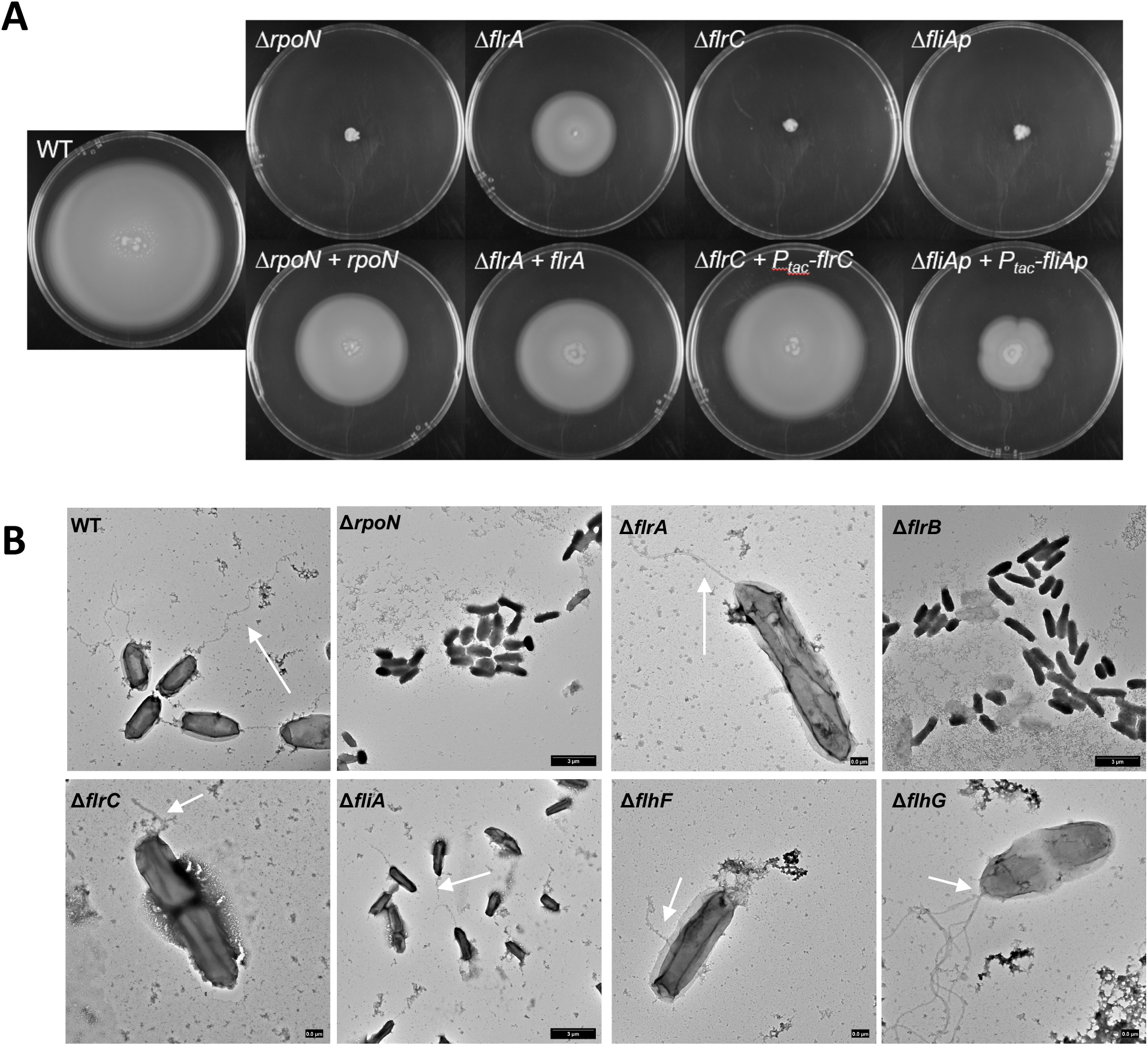
Swimming and flagellar phenotypes of *V. campbellii* flagellar regulator mutants. Soft agar (0.3%) swim plates showing swim halo phenotypes for wild-type and mutant DS40M4 strains. Top panel shows deletion mutants. Bottom panel shows deletion mutants that were complemented at an ectopic locus (*luxB*). B) Transmission electron microscopy of wild- type and mutant DS40M4 strains, grown in LM liquid culture. White arrows point to flagellar protrusions in strains.

However, the Δ*flrA* mutant demonstrates swimming motility, albeit partially reduced compared to the wild-type strain. We complemented each mutant at an ectopic locus in the genome. For *rpoN* and *flrA*, our TSS analyses indicated that these genes are likely the first gene in the operon, thus we complemented these using their cognate promoter. However, for *flrC* and *fliA*, it was unclear from our dRNA-seq data which promoter(s) controls expression of these genes, and thus we complemented *flrC* and *fliA* using an IPTG-inducible promoter. In each case, complementation results in increased levels of motility compared to the deletion strain (Fig. 4A).

We also directly observed the presence or absence of flagella via transmission electron microscopy (TEM) to compare the structural effects of deletion of flagellar regulators (Fig. 4B). We chose to assess deletion strains for each of the four flagellar transcriptional regulators (Δ*rpoN*, Δ*flrA*, Δ*flrC*, and Δ*fliA*) as well as Δ*flrB*, Δ*flhF*, and Δ*flhG*, which act within the four-tiered transcriptional hierarchy and have observable flagellar patterns in other vibrios (41, 47, 48). When *rpoN* and *flrB* are deleted, no flagella are detectable via TEM. The Δ*flrC* cells are also largely aflagellate, however, some observable short cable structures did protrude from some of the cells (Fig. 4B). When *flhF* is deleted, cells were largely aflagellate except for a few that produce partial and/or nonpolar flagella. Conversely, Δ*flhG* mutants demonstrate hyperflagellation. These findings correlate with prior studies in *Vibrio* species indicating FlhF’s role in flagellar localization and FlhG’s role in regulating flagellar number (41, 47). Deletion of either *flrA* or *fliA* had no observable impact on detectable flagella via TEM (Fig. 4B). In the Δ*fliA* mutant this may be explained by the role of FliA in primarily regulating chemotaxis and motor genes rather than structural genes, though FliA is typically required for expression of alternate flagellins in vibrios (19). However, it is notable that the Δ*flrA* mutant produces intact flagella, conflicting with the models in other *Vibrio* species in which it is one of the earliest required transcriptional regulators for expression of flagellar genes (18, 19). Altogether, these results largely support previous findings in other vibrios with one major exception: we conclude that FlrA is not required for flagellar production or motility in *V. campbellii*.

### *The lateral flagellar gene* lafK *does not compensate for absence of* flrA

In addition to encoding the polar flagellar gene system, *V. campbellii* also encodes a set of genes homologous to known *V. parahaemolyticus* lateral flagellar genes (26). The lateral flagellar gene system is a separate set of similar but distinct genes which produce numerous, non-sheathed lateral flagella and which are encoded in Chromosome II rather than I. Historically, lateral flagella have been characterized in *V. parahaemolyticus*, but other vibrios such as *V. alginolyticus* and *V. campbellii* have been found to either produce or encode these genes as well (26, 30, 31, 49). Importantly, the lateral flagellar σ^54^-dependent transcriptional regulator LafK cross-regulates Class II polar flagellar genes in the absence of the FlrA homolog (FlaK) in *V. parahaemolyticus* (46). *V. campbellii* encodes a *lafK* homolog in its genome. We hypothesized that *V. campbellii* LafK regulation facilitates the partially motile phenotype in the Δ*flrA* mutant strain. The *V. campbellii* Δ*lafK* mutant demonstrates no change in swimming motility compared to wild-type, mimicking the results in *V. parahaemolyticus* (Fig. 5). However, unlike *V. parahaemolyticus*, the Δ*lafK* Δ*flrA* mutant shows no further decrease in motility compared to the Δ*flrA* mutant (Fig. 5). Thus, we conclude that *V. campbellii* LafK does not compensate for FlrA.

**Figure 5.**
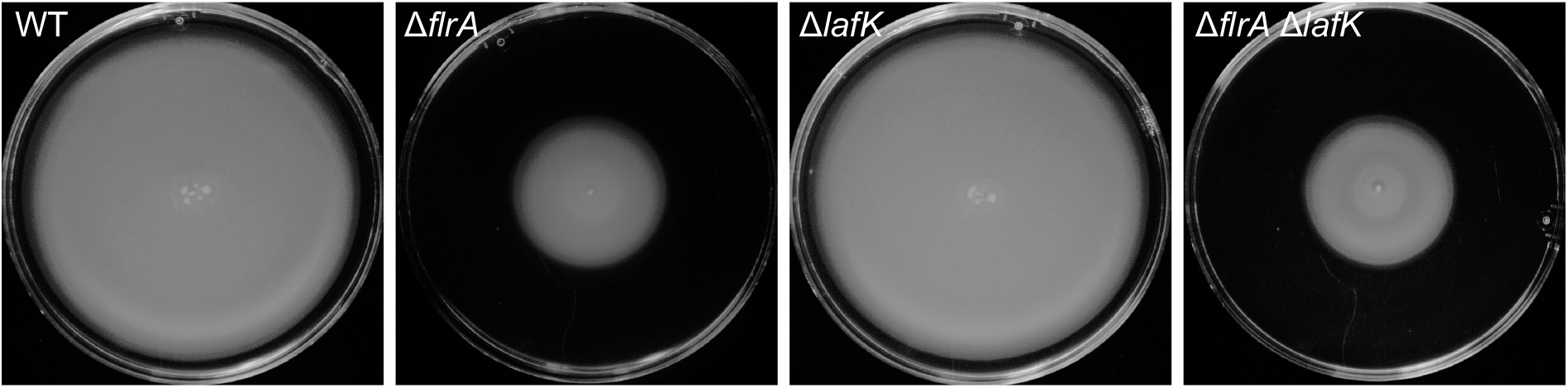
LafK does not regulate swimming motility in *V. campbellii*. Soft agar (0.3%) swim plates showing swim halo phenotypes for wild-type and mutant strains of DS40M4.

Due to the lack of observed LafK-mediated polar flagellar regulation, we hypothesized that *V. campbellii* may not have an active lateral flagellar system for swarming motility. Although *V. campbellii* encodes lateral flagella, few studies have characterized these flagella or the capacity for swarming motility in *V. campbellii* (30, 31). We tested to see whether *V. campbellii* was capable of swarming motility by performing swarm plate assays with varying agar concentrations. *V. parahaemolyticus* is capable of swarming at agar concentrations of 1.5%, while other swarming bacteria such as *B. subtilis* swarm at agar concentrations between 0.3- 1.0% (50). Under each condition tested, we were unable to observe swarming in *V. campbellii* DS40M4 (Fig. S1). We also assayed for swarming motility in three additional *V. campbellii* strains but no swarming was observed, whereas we were able to visualize swarming in *V. parahaemolyticus* as a positive control (Fig. S1). Together, our data suggest that despite encoding lateral flagellar genes, *V. campbellii* DS40M4 has an inactive lateral flagellar system under the conditions we tested. In addition, we conclude that the lateral flagellar regulator LafK does not compensate for FlrA activity in a Δ*flrA* mutant.

### Transcriptome profiling of flagellar regulators

To identify the *V. campbellii* genes that are controlled by the transcriptional regulators σ^54^, FlrA, FlrC, and FliA, we compared the transcription profiles between wild-type and isogenic mutant strains lacking one of these four genes (*rpoN, flrA, flrC*, and *fliA*, respectively) using RNA-seq (Dataset S1). Our goal was to sort genes into different classes in the regulatory hierarchy based on gene expression relative to wild-type similar to previous studies (16). We predicted that genes expressed earlier in the transcriptional hierarchy are only regulated by the earliest flagellar regulator(s) and not any downstream regulators, whereas genes expressed later in the hierarchy will be affected by all upstream regulators.

Figure 6 shows the expression profiles of flagellar genes for Δ*rpoN*, Δ*flrA*, Δ*flrC*, and Δ*fliA* compared to wild-type determined by RNA-seq (Fig. 6; differential gene expression data are presented in Dataset S1). These data revealed that much of the downstream hierarchy is organized similarly to the transcriptional hierarchy established in other vibrios (16, 27).

**Figure 6.**
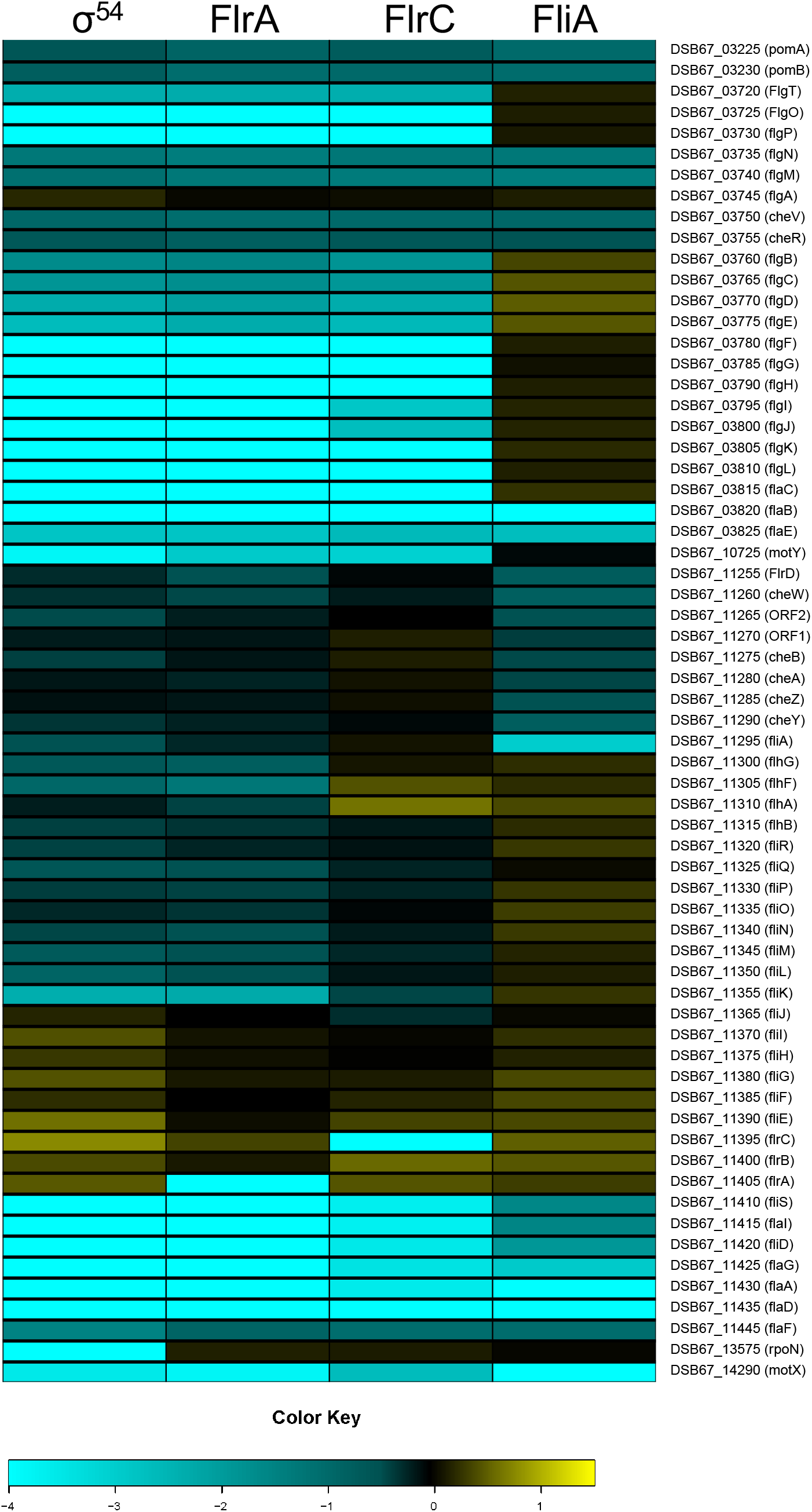
Expression patterns of Class I-III polar flagellar genes. Cluster analysis of RNA-seq data for polar flagellar genes showing expression in each mutant strain compared to wild-type. Flagellar gene names and locus tags are indicated. The log_2_ fold- change in gene expression relative to wild-type is represented by the color key.

Specifically, FliA-dependent expression patterns are very similar to Class IV flagellar genes in other vibrios, where we observed that transcription decreases in the absence of FliA and/or upstream regulators for *pomAB, flgMN, cheVR, flaBE, flrD, cheW*ORF1ORF2, *cheBAZY, fliSflaIfliDflaGflaA, flaDF*, and *motX*. Similarly, FlrC-dependent expression patterns are comparable to Class III in other vibrios as well, as we identified several genes clusters that were decreased in expression in the absence of FlrC: *flgT, flgOP, flgBCDE, flgFGHIJ, flgKLflaC, fliKLMNOPQR, flhB*, and *motY*.

The expression patterns for FlrA showed the largest deviation from studies of other vibrios. In *V. cholerae*, FlrA is required for regulation of several flagellar operons, including *fliEFGHIJ, flhAFG, flrBC*, and *fliA* (16, 19). Deletion of *flrA* also results in decreased expression of downstream genes controlled by either FlrC or FliA, such as hook, rod, chemotaxis, and flagellin genes Conversely, our data show that FlrA is not required for expression of *fliEFGHIJ* or *flrBC*, which are expressed at levels similar to wild-type in the absence of either FlrA or σ^54^ (Fig. 6, Dataset S1). The only unique set of genes solely affected by Δ*flrA* are *flhAFG*. This result correlates with the swim plate results in which *flrA* is not required for swimming motility, but deletion of *flrA* reduces swim halo size. Notably, however, *fliK, fliLMNOPQR* and *flhB* expression patterns are decreased in both a Δ*flrA* mutant than in a Δ*flrC* mutant, which could suggest that both FlrA and FlrC regulate these genes. From these data, we conclude that FlrA-dependent regulatory patterns in *V. campbellii* DS40M4 is distinct from that of FlrA in *V. cholerae*. In addition, transcription of genes classified as Class II in *V. cholerae* – *fliEFGHIJ, flrA*, and *flrBC* – is independent of the four regulators studied here in *V. campbellii*.

### FlrA and FlrC have both separate and overlapping regulons

The results of our RNA-seq analysis indicate that some flagellar genes are decreased in expression in both the Δ*flrA* and Δ*flrC* backgrounds compared to wild-type. FlrA and FlrC are both bacterial enhancer binding proteins (bEBPs) that share 39.79% identity and serve as σ^54^- dependent transcriptional regulators (44, 51, 52). Thus, we hypothesized that FlrA shares a co-regulon with FlrC, in addition to their distinct regulons of genes. To test this, we performed qPCR on wild-type, Δ*flrA*, Δ*flrC*, and Δ*flrA* Δ*flrC* strains. In addition, we complemented *flrA* and *flrC* in the double Δ*flrA* Δ*flrC* mutant background. We chose to examine genes from each class based on our RNA-seq data: *flhF* is regulated by FlrA, *flgO* is regulated by FlrC, and *fliK* is co-regulated by FlrA and FlrC. As a control, we assessed gene expression of *flaA*, a gene controlled by FliA late in the hierarchy, thus affected by all upstream regulators. Similar to our RNA-seq results, we find that *flhF* is more strongly decreased in expression in Δ*flrA*, whereas *flgO* is more strongly decreased in in Δ*flrC* compared to wild-type (Fig. 7A, 7B). However, deletion of Δ*flrA* and Δ*flrC* have a significant effect on both genes. In addition, deletion of either *flrA* or *flrC* partially decreases in expression of *fliK* compared to wild-type, whereas the deletion of both *flrA* and *flrC* further decreases *fliK* expression compared to the single mutants (Fig. 7C). Complementation of *flrA* or *flrC* in the Δ*flrA* Δ*flrC* background restores *flgO* levels to the single-mutant level (Fig. 7B), whereas complementation did not affect *flhF* or *fliK* levels (Fig. 7A, 7B). As predicted, we also find that deletion of either *flrA* or *flrC* significantly impairs *flaA* expression (Fig. 7D). We observe additive effects on regulation of *fliK* and *flgO*, suggesting that both FlrA and FlrC control expression of the *fliK* and *flgO* operons. Although deletion of *flrC* does significantly decrease *flhF* expression, the deletion of *flrA* and *flrC* is not additive, suggesting that FlrA is the dominant regulator (Fig. 7A). From these data, we conclude that FlrA and FlrC have compensatory roles and co-regulate most of the Class II genes in the flagellar hierarchy.

**Figure 7.**
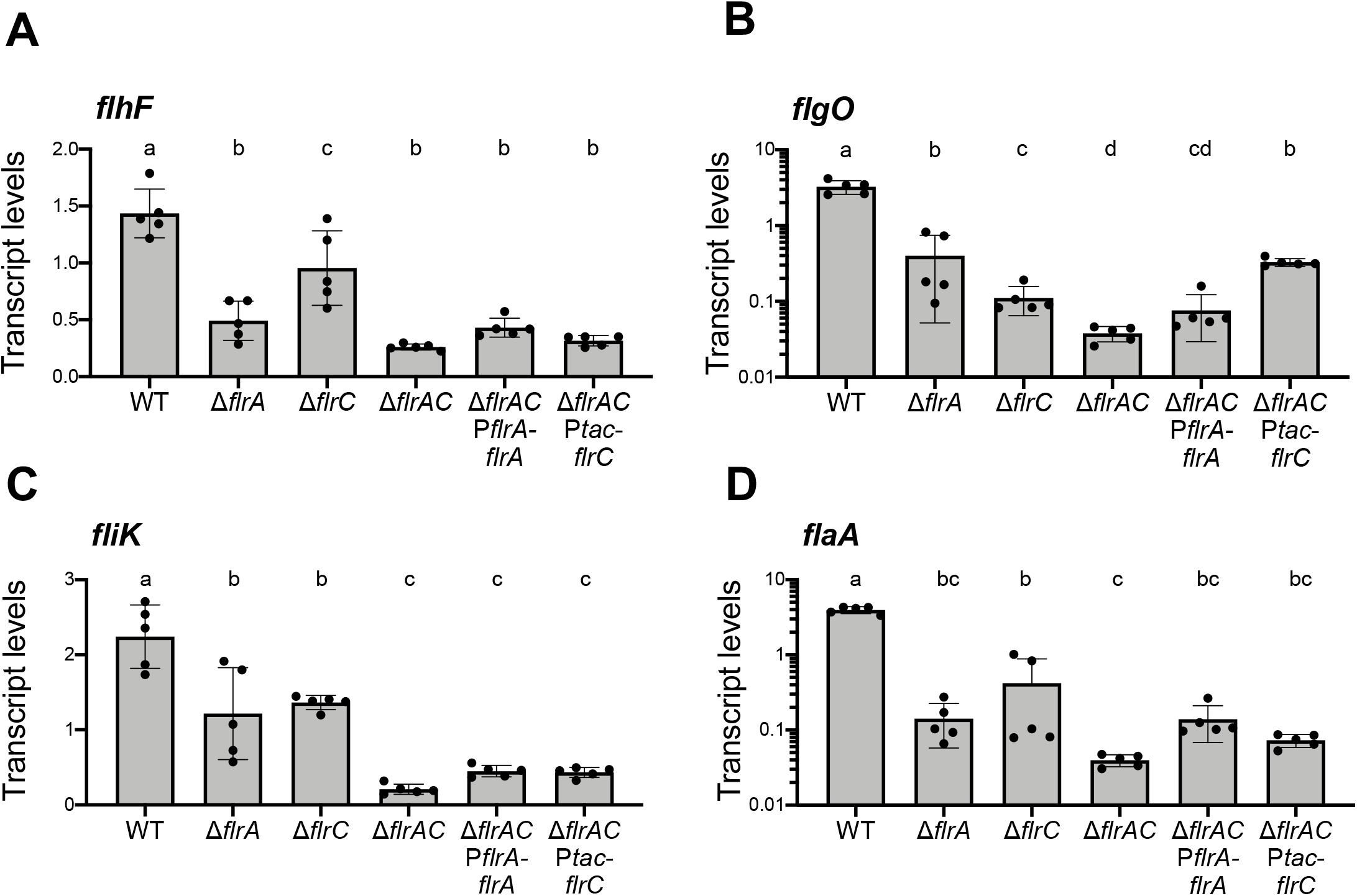
FlrA and FlrC have separate and overlapping regulons. Data shown are absolute transcripts quantified by qPCR of genes from representative operons *flhF*, b) *flgO*, c) *fliK*, d) *flaA* regulated by FlrA and FlrC in the wild-type, Δ*flrA*, Δ*flrC*, Δ*flrA* Δ*flrC*, Δ*flrA* Δ*flrC* P_*flrA*_-*flrA*, Δ*flrA* Δ*flrC* P_*tac*_-*flrC* strains. Different letters indicate significant differences in 2-way analysis of variance (ANOVA) of log-transformed data followed by Tukey’s multiple comparison’s test (*n* = 5, *p* < 0.05).

## Discussion

Swimming motility is employed by many different *Vibrio* species for taxis toward or away from stimuli. The genetic and regulatory networks underlying flagellar swimming motility have primarily been studied in *V. cholerae, V. parahaemolyticus*, and *V. fischeri* by testing individual flagellar mutants or by impairing specific flagellar gene intervals (16, 22, 23, 45). By exploiting the power of natural transformation, we were able to efficiently delete every known gene in the five flagellar and chemotaxis genetic loci in *V. campbellii*. Our work here represents both the first broad examination of the polar flagellar genetic network in *V. campbellii*, as well as one of the first studies to characterize the complete set of known genes in the flagellar and chemotaxis genomic regions in a *Vibrio* species using reverse genetics.

Of the 64 genes we examined in this study, the phenotypes of most mutants are consistent with predicted outcomes based on prior studies in other bacterial species. The majority of mutants we characterized are semimotile, as have been observed in previous flagellar systems as well, suggesting that these strains are still capable of synthesizing a partially functional flagellum (22). These strains include those impaired in components of the basal body or the outer flagellar rings (L-ring, P-ring, H-ring), though notably hook and rod mutants were not observed in this mutant class. We found that flagellar hook and rod mutants were nonmotile, likely because these structures are critical for downstream flagellar assembly steps or flagellar joint functionality, which aligns with previous findings in other bacteria (22, 53). We were surprised to find that flagellar filament cap mutants lacking either *fliT* or *fliD* demonstrated detectable motility, though this result does support previous findings in *V. parahaemolyticus* in which absence of *fliD* does not completely inhibit motility unlike in *S. enterica* (54, 55). Most of the nonmotile mutants we identified were also aflagellate, except for the motor mutants, due to the stepwise nature of assembly of the flagellum in bacteria. Notably, prior studies that characterized presence or absence of a flagellum did so using western immunoblots to measure the presence of polar flagellin production (22). Rather, in this study we directly observed the presence or absence of a flagellum using fluorescence staining and microscopy. However, this approach excludes cells that continue to produce and secrete downstream flagellar components (such as flagellins), regardless of whether these secreted products correlate with an intact flagellum. Both approaches have their advantages, and this distinction could account for differences we observed between flagellate and aflagellate mutants in this study versus previous work.

Importantly we found that none of the flagellin genes (*flaABCDEF*) are absolutely required for motility in *V. campbellii*. Only deletion of *flaC* impairs motility compared to the other mutants, which all resemble wild-type. FlaC is most closely related to FlaA in *V. cholerae*, which is the only filament protein required and sufficient for motility (56). The reduced motility in the *V. campbellii* Δ*flaC* mutant correlates with most other vibrios that also encode a predominant flagellin subunit that impairs (but does not eliminate) motility when deleted (54, 57–59).

A few *Vibrio* species such as *V. parahaemolyticus, V. alginolyticus*, and *V. campbellii* also encode lateral flagellar genes in addition to their polar flagellar system, which provide them the capacity for swarming motility over surfaces (26, 30, 31). While characterizing the polar flagellar system in *V. campbellii* in this study, we also studied the lateral flagella and swarming motility in an attempt to describe this behavior as well. When comparing several different strains of *V. campbellii* using different hard agar concentrations, we were unable to observe swarming motility in any of these strains. We also did not observe additional flagella beyond the presence of the polar flagellum using TEM of cells prepared from hard agar or soft agar plates in the BB120 strain (Fig. 4B, data not shown). Notably, the first study to identify lateral flagella in *V. campbellii* (then called *Beneckea campbellii*) observed that only some strains produced both polar and lateral flagella, while many of the strains only produced a single polar flagellum (31). Taking our observations here and the results found in previous studies of *V. campbellii*, we believe it is likely that the *V. campbellii* strains we examined have an inactive lateral flagellar system under our laboratory conditions.

Our transcriptome analysis of the flagellar regulatory mutants in *V. campbellii* highlighted some unique features of the flagellar regulatory hierarchy, which allowed us to construct a model for the flagellar regulatory hierarchy in *V. campbellii* (Fig. 8). First, we note that while FlrA plays an observable role in the flagellar regulatory hierarchy, it is also not necessary for *V. campbellii* cells to be motile on swim plates. Genes that were classified as Class II in *V. cholerae* are mostly expressed independently of both σ^54^ and FlrA in *V. campbellii*, except for *flhAFG* and *fliA* (16). This presents two possibilities for the flagellar regulatory hierarchy in *V. campbellii*. One possibility is that the four-tiered flagellar hierarchy is actually a three-tiered hierarchy in *V. campbellii*, where some previously identified Class II genes such as *fliEFGHIJ* and the downstream flagellar regulators like *flrB* and *flrC* are expressed constitutively and independently of other regulators like σ^54^. Taking the data we have presented here into account, without bias from previously defined hierarchies from other vibrio flagellar species, this is our proposed model for flagellar transcriptional regulation in *V. campbellii* (Figure 8). Alternatively, however, without a stepwise checkpoint system in place if multiple Class I operons were expressed constitutively in *V. campbellii*, this could cause a competitive disadvantage because cells would needlessly waste resources in expressing flagellar gene products. Previous reports show that flagellar synthesis is a costly behavior in bacteria, and expression of flagellar genes can impair growth (60–62). An alternative hypothesis is that an unidentified regulator controls the expression of these independently expressed Class I genes in *V. campbellii*. While this model would preserve the four-tiered stepwise mechanism of flagellar regulation conserved by other vibrios, our current data suggest the Class I genes are independent of σ^54^ regulation and we have not found any evidence of a novel regulator for these genes in *V. campbellii*. Thus the first hypothesis is our proposed model that we present in Figure 8. We anticipate that future work will uncover the presence or absence of a potential novel Class I flagellar regulator.

**Figure 8.**
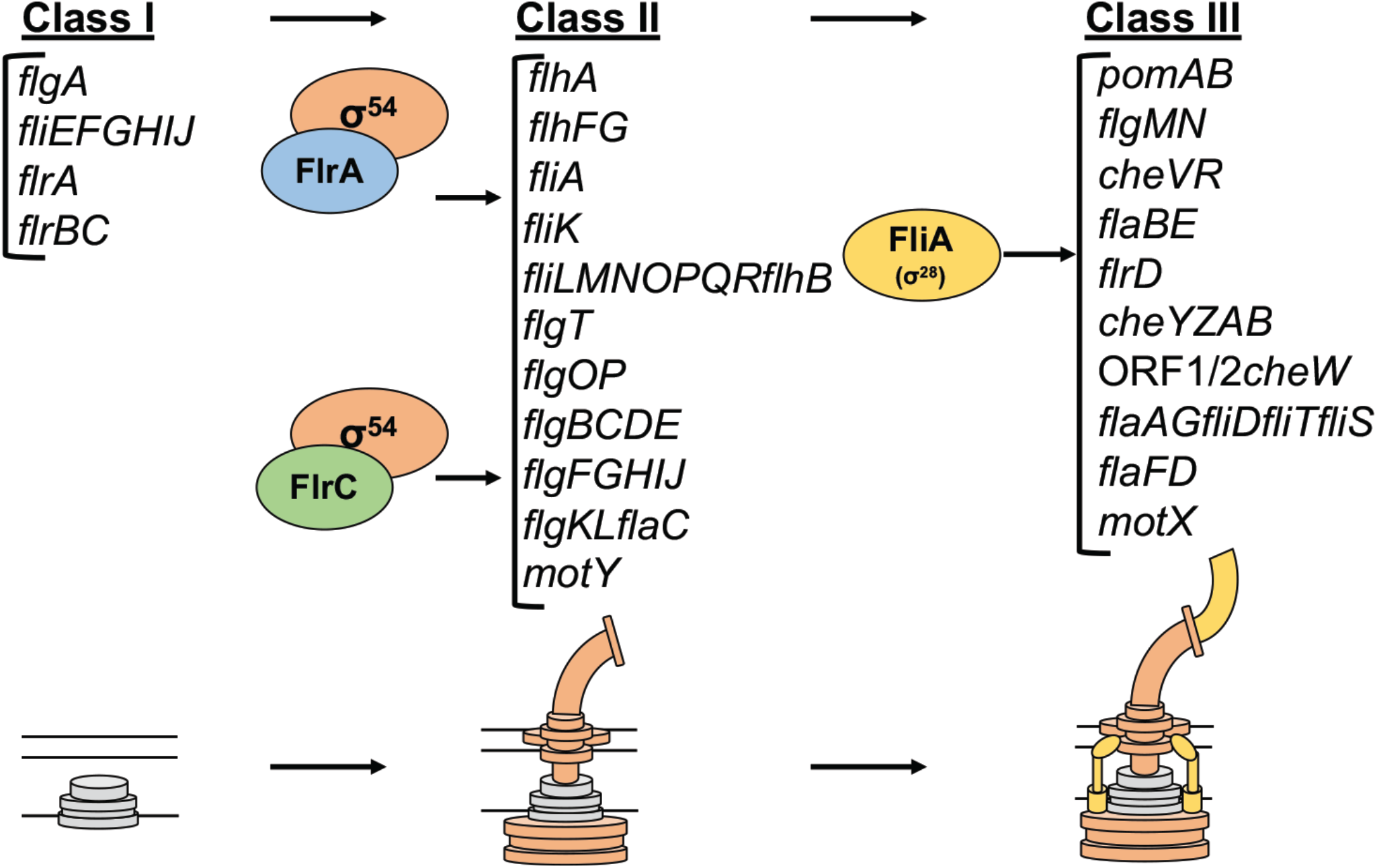
Transcriptional hierarchy of polar flagellar genes in *V. campbellii*. Model of transcriptional hierarchy for polar flagellar genes in *V. campbellii* DS40M4. Class I encodes early structural components of the flagella as well as the Class II regulators. Class II genes depend on regulation by σ^54^ and one or both σ^54^-dependent transcriptional activators, FlrA and FlrC. These genes include most structural components of the flagellum, including the export apparatus, hook, rod, ATPase, flagellin, flagellar rings, and more. Finally, FliA (σ28) activates Class III genes, which encode additional flagellins, chemotaxis proteins, and the motor/stator.

Another critical difference in our proposed hierarchy model is how Class II genes are regulated in *V. campbellii*. In our RNA-seq analysis, FlrC in *V. campbellii* showed very similar expression profiles to those observed in *V. cholerae*, thus our model groups Class II_*V*.*camp*_ and Class III_*V*.*chol*_ genes very similarly between both species (16). However, as we noted earlier, FlrA still plays an observable role in regulation of downstream flagellar genes such as Class II and Class III, and we found that *fliKLMNOPQR* and *flhB* showed stronger decreases in expression in a Δ*flrA* compared to a Δ*flrC* mutant. When testing these further with qPCR in both single mutants and a Δ*flrA*Δ*flrC* double mutant, we found that deletion of either FlrA or FlrC significantly impaired expression of genes from the opposite regulon. In addition, a subset of genes classified as FlrC-dependent in both our *V. campbellii* RNA-seq data and previous *V. cholerae* models (17) showed additive downregulation when both σ^54^-dependent flagellar regulators were deleted. Based on these findings, we propose that FlrA and FlrC are both partially interchangeable σ^54^-dependent transcriptional regulators, and that certain Class II flagellar genes are either regulated by one of the two regulators or co-regulated by both FlrA and FlrC. Future work will more carefully tease apart this class of regulation to better understand the mechanisms behind regulation by both bEBPs. Finally, while Class IV gene regulation in *V. cholerae* is very similar to our findings for Class III in *V. campbellii*, we noted that *fliS, flaI, fliD, flag, flaA* and ORF1/ORF2/*flrD* display FliA-dependent expression patterns in *V. campbellii*, thus grouping them with other Class III genes in the regulatory hierarchy (16).

Although the flagellar genes in *V. campbellii* are conserved and share similar chromosomal organization with other *Vibrio* species, we have shown that the regulation of the polar flagellum in *V. campbellii* is distinct in several ways. Furthering our understanding of the regulation of motility in different *Vibrio* species is important because motility and taxis in response to external signals are key behaviors of bacteria in various niches. Together, our work here has established tools and broadened our understanding of the flagellar gene network in *V. campbellii* that will be valuable in future work both in studying swimming motility in *V. campbellii* and comparing flagellar regulation across the *Vibrionaceae*.

## Experimental procedures

### Strains and growth conditions

All bacterial strains and plasmids are listed in Tables S1 and S2. *V. campbellii* strains were grown at 30°C on Luria marine (LM) medium (Lysogeny broth supplemented with an additional 10 g NaCl per L). Instant ocean water (IOW) medium was used in chitin-independent transformations; it consists of Instant Ocean sea salts (Aquarium Systems, Inc.) diluted in sterile water (2X = 28 g/l). Transformations were outgrown in LBv2 (Lysogeny Broth medium supplemented with additional 200 mM NaCl, 23.14 mM MgCl_2_, and 4.2 mM KCl). When necessary, strains were supplemented with kanamycin (100 μg/ml), spectinomycin (200 μg/ml), or trimethoprim (10 μg/ml).

### Analysis of Vibrio flagellar genes

A dataset comprising of all the protein sequences from the various *Vibrio* species (including DS40M4) was clustered into groups based on sequence similarity using cd-hit version v4.8.1-2019-0228 (63) (parameters: -M 0 -g 1 -s 0.8 -c 0.4 -n 2 -d 500). The protein sequences from the polar flagellar gene clusters were compared against the corresponding DS40M4 flagellar gene using BLASTp and the percent identity of the best scoring homolog from each *Vibrio* species was recorded and plotted in the heatmap. The six flagellins overlapped in a single cluster due to high sequence identity and were resolved manually.

### Primer design and strain construction

A candidate primer set was generated for deletion of each gene consisting of two primer pairs chosen to maximize the portion of the gene removed such that (1) any overlapping genes were left complete and (2) all four primers met our criteria for validity. A valid primer is between 25 and 30 basepairs with a preference for shorter primers; contains no homopolymers of 5 or more bases; ends in cytosine or guanine; and has a T_m_ between 58 and 63 degrees as determined by the BioPerl implementation of Allawai and SantaLucia, 1997. Custom scripts are available on GitHub (https://github.com/Juliacvk/vibrio_primer_design).

All primers are listed in Table S3. These primers were used to generate linear transforming DNAs (tDNA) by splicing-by-overlap extension (SOE) PCR as previously described (34). Briefly, an UP arm containing ∼3kb homology of the area upstream of the gene to be deleted was PCR amplified via a F1 and R1 primer set. Similarly, a DOWN arm containing ∼3kb homology of the area downstream of the gene to be deleted was PCR amplified via a F2 and R2 primer set. For unselected tDNA products, the UP and DOWN arms were used as template and amplified via the F1 and R2 primers to make a SOE product that lacked an antibiotic marker. Unselected tDNA products were cotransformed with a selected tDNA product, which targets the nonfunctional *luxB* site in DS40M4 and inserts an antibiotic resistance cassette (for either spectinomycin or trimethoprim). These tDNAs were used to construct *V. campbellii* DS40M4 strains using previously published methods (64). Briefly, DS40M4 were grown overnight in LBv2 at 30°C, shaking, with antibiotic and 100 μM IPTG. The next day the culture was back-diluted into IOW containing 100 μM IPTG and tDNA. The tDNA marked with an antibiotic resistance cassette was added and cells were incubated statically at 30°C for 4-6 hours. LBv2 was then added and the cells were incubated shaking at 30°C for an additional 1-2 hours before streaking cultures out on LM plates with respective selective antibiotics. Following natural transformation, strains containing the correct target mutation were identified via colony PCR with a forward and reverse detection primer and further confirmed by sending the products to Eurofins Scientific for sequencing.

### Motility assays

Swimming motility was measured by performing swimming assays on soft agar (media containing 0.3% agar). *V. campbellii* strains were grown overnight in LM media, and were diluted to OD_600_ = 0.5 the following day. Five µl of the diluted cultures were stabbed and pipetted into the center of soft agar LM plates and grown in a humid container at 30°C for 24 hours. After growth, plates were imaged via AlphaImager HP Imaging System (ProteinSimple) and compared for size of swim halos.

Swarm assays were performed using LM hard agar plates (0.7%, 1.0%, and 1.5% agar). *V. campbellii* strains were grown overnight in LM media, and were concentrated to OD_600_ = 10 the following day. Five µl of the cultures were spotted onto the surface of LM swarm plates and grown in a humid container at 30°C for 24 hours. After growth, plates were measured for radius of swarming.

### RNA extraction, qRT-PCR, and RNA-seq

Strains were inoculated in 5 ml LM and grown overnight shaking at 30°C at 275 RPM. Each strain was back-diluted 1:1,000 in 5 ml LM and grown shaking at 30°C at 275 RPM until they reached an OD_600_ = ∼0.5. Cells were collected by centrifugation at 3,700 RPM at 4°C for 10 min, the supernatant was removed, and the cell pellets were flash frozen in liquid N_2_ and stored at -80°C. RNA was isolated from pellets using a TRIzol/chloroform extraction protocol as described (65) and treated with DNase via the DNA-free™ DNA Removal Kit (Invitrogen). RNA- seq was performed as described previously (66). Sequence data were deposited in the National Center for Biotechnology Information Gene Expression Omnibus with accession number GSE167483.

Real-time qRT-PCR (quantitative reverse transcription PCR) was used to quantify transcript levels using the SensiFast SYBR Hi-ROX One-Step Kit (Bioline) according to the manufacturer’s guidelines. 1 µg of each RNA template was converted to cDNA using SuperScript IV reverse transcriptase (Invitrogen). All reactions were then performed using a LightCycler 4800 II (Roche) with 0.4 µM of each primer and 5 ng of template RNA (10 µl total volume). Primers were designed to have the following parameters: amplicon size of 100 bp, primer size of 20-28 nt, and melting temperature of 55-60°C.All qPCR experiments were normalized to *hfq* expression and were performed with 5 biological replicates and 2 technical replicates.

### dRNA-Seq analysis

cDNA libraries from RNA (isolated as described above) were constructed as described previously (67) by Vertis Biotechnology AG (Freising, Germany) and sequenced using an Illumina NextSeq 500 machine in single-read mode (75 bp read length). The raw, demultiplexed reads and coverage files have been deposited in the National Center for Biotechnology Information Gene Expression Omnibus with accession code GSE147616.

### Transcriptional start site analysis

Transcriptional start site (TSS) prediction was performed using the program TSSpredator on normalized coverage data (Dataset S2). TSS classified as Primary or Secondary are the ones located upstream of a gene, not further apart than the chosen UTR length. Of those, the Primary TSS is the one with the strongest expression. The ones within the annotated gene located on the sense strand are classified as Internal, while the ones which are on the antisense strand not farther than the chosen UTR length are classified as Antisense. Finally, those which are not within the vicinity of any of the annotated genes are classified as Orphans. The TSS data are presented in Dataset S2.

### Flagellar labelling and microscopy

Fluorescence microscopy was performed with a Nikon Eclipse 80i microscope equipped with a Plan Apo 100× phase-contrast objective. Cells were grown in LM media to OD_600_ ∼0.5, near mid-log phase, and 1 ml of culture volume was concentrated by centrifugation at 4,000 x g for one minute at room temperature and resuspended in 50 µl phosphate buffer saline (PBS). 1µl NanoOrange dye (ThermoFischer Scientific) was mixed with the cells and given 15 min for staining to occur. NanoOrange-stained cells and flagella were visualized with a FITC HYQ filter cube (excitation filter, 460 to 500 nm; barrier filter, >590 nm). Images were captured with a Photometrics Coolsnap HQ2 camera and processed using MetaMorph 7.7.9.0 image software (Universal Imaging Corp.).

For TEM, strains were inoculated in 5 ml LM and grown overnight shaking at 30°C at 275 RPM. Each strain was back-diluted 1:1,000 in 5 ml LM and grown shaking at 30°C at 275 RPM until they reached OD_600_ ∼0.5, near mid-log phase. Cells were collected by centrifugation at 4,000 x g for one minute at room temperature and washed with phosphate buffer saline (PBS). Cells were collected on grids and stained in 2% uranyl acetate. The grids were commercially prepared continuous carbon films on 400 mesh copper grids obtained from Ted Pella, Inc. Electron microscopy was performed at nominal magnifications ranging from 2500x to 50,000x using a JEOL JEM 14000plus TEM operating at 120kV. Images were recorded with a Gatan OneView camera using 0.5 s exposures and Gatan’s buit-in motion correction.

## Acknowledgments

We thank Victoria Lydick for providing excellent technical support in the lab. We also thank David Gene Morgan and the IU Electron Microscopy Center for their help with imaging vibrio flagella with TEM. Finally, we would like to thank Daniel Kearns for project guidance and comments on the manuscript.

## Funding

This work was supported by the National Institutes of Health grant R35GM124698 to JVK.

## Notes

### Competing Interest Statement

The authors have declared no competing interest.

## References

1. Boin MA, Austin MJ, Häse CC. 2004. Chemotaxis in Vibrio cholerae. FEMS Microbiol Lett 239:1–8.

2. Ottemann KM, Miller JF. 1997. Roles for motility in bacterial – host interactions. Mol Microbiol 24:1109–1117.

3. Norsworthy AN, Visick KL. 2013. Gimme shelter: how Vibrio fischeri successfully navigates an animal’s multiple environments. Front Microbiol 4:1–14.

4. Reidl J, Klose KE. 2002. Vibrio cholerae and cholera: Out of the water and into the host. FEMS Microbiol Rev 26:125–139.

5. Letchumanan V, Chan KG, Lee LH. 2014. Vibrio parahaemolyticus: A review on the pathogenesis, prevalence, and advance molecular identification techniques. Front Microbiol 5:1–13.

6. Pujalte MJ, Sitjà-Bobadilla A, Macián MC, Belloch C, Álvarez-Pellitero P, Pérez-Sánchez J, Uruburu F, Garay E. 2003. Virulence and molecular typing of Vibrio harveyi strains isolated from cultured dentex, gilthead sea bream and European sea bass. Syst Appl Microbiol 26:284–292.

7. Ruby EG. 1996. Lessons from a cooperative, bacterial-animal association: The Vibrio fischeri-Euprymna scolopes light organ symbiosis. Annu Rev Microbiol 50:591–624.

8. Austin B, Zhang XH. 2006. Vibrio harveyi: A significant pathogen of marine vertebrates and invertebrates. Lett Appl Microbiol 43:119–124.

9. Zhang X-H, He X, Austin B. 2020. Vibrio harveyi: a serious pathogen of fish and invertebrates in mariculture. Mar Life Sci Technol 2:231–245.

10. Haldar S, Chatterjee S, Sugimoto N, Das S, Chowdhury N, Hinenoya A, Asakura M, Yamasaki S. 2011. Identification of Vibrio campbellii isolated from diseased farm-shrimps from south India and establishment of its pathogenic potential in an Artemia model. Microbiology 157:179–188.

11. Daniels NA, Mackinnon L, Bishop R, Altekruse S, Ray B, Hammond RM, Thompson S, Wilson S, Bean NH, Griffin PM, Slutsker L. 2000. Vibrio parahaemolyticus infections in the United States, 1973-1998. J Infect Dis 181:1661–1666.

12. Lee JH, Rho JB, Park KJ, Kim CB, Han YS, Choi SH, Lee KH, Park SJ. 2004. Role of flagellum and motility in pathogenesis of Vibrio vulnificus. Infect Immun 72:4905–4910.

13. Yang Q, Defoirdt T. 2015. Quorum sensing positively regulates flagellar motility in pathogenic Vibrio harveyi. Environ Microbiol 17:960–968.

14. Rugel AR, Klose KE. 2011. Vibrio cholerae Flagellar Synthesis and Virulence. Epidemiol Mol Asp Cholera 203–212.

15. Zhu S, Kojima S, Homma M. 2013. Structure, gene regulation and environmental response of flagella in Vibrio. Front Microbiol 4:1–9.

16. Syed KA, Beyhan S, Correa N, Queen J, Liu J, Peng F, Satchell KJF, Yildiz F, Klose KE. 2009. The Vibrio cholerae flagellar regulatory hierarchy controls expression of virulence factors. J Bacteriol 191:6555–6570.

17. Echazarreta MA, Klose KE. 2019. Vibrio Flagellar Synthesis 9:1–11.

18. Millikan DS, Ruby EG. 2003. FlrA, a σ54-dependent transcriptional activator in Vibrio fischeri, is required for motility and symbiotic light-organ colonization. J Bacteriol 185:3547–3557.

19. Prouty MG, Correa NE, Klose KE. 2001. The novel σ54- and σ28-dependent flagellar gene transcription hierarchy of Vibrio cholerae. Mol Microbiol 39:1595–1609.

20. Burnham PM, Kolar WP, Hendrixson DR. 2020. A polar flagellar transcriptional program mediated by diverse two-component signal transduction systems and basal flagellar proteins is broadly conserved in polar flagellates. MBio 11:1–25.

21. Correa NE, Barker JR, Klose KE. 2004. The vibrio cholerae FlgM homologue is an anti-σ28 factor that is secreted through the sheathed polar flagellum. J Bacteriol 186:4613– 4619.

22. Kim Y, Carter Llmc. 2000. Analysis of the Polar Flagellar Gene System of Vibrio parahaemolyticus. J Bacteriol 182:3693–3704.

23. Brennan CA, Mandel MJ, Gyllborg MC, Thomasgard KA, Ruby EG. 2013. Genetic determinants of swimming motility in the squid light-organ symbiont Vibrio fischeri. Microbiologyopen 2:576–594.

24. Millikan DS, Ruby EG. 2002. Alterations in Vibrio fischeri motility correlate with a delay in symbiosis initiation and are associated with additional symbiotic colonization defects. Appl Environ Microbiol 68:2519–2528.

25. McCarter LL. 2006. Motility and Chemotaxis, p. 113–132. In Thompson, FL, Austin, B, Swings, J (eds.), The Biology of Vibrios. ASM Press, Washington, DC, USA.

26. McCarter LL. 2004. Dual flagellar systems enable motility under different circumstances. J Mol Microbiol Biotechnol 7:18–29.

27. Gode-Potratz CJ, Kustusch RJ, Breheny PJ, Weiss DS, McCarter LL. 2011. Surface sensing in Vibrio parahaemolyticus triggers a programme of gene expression that promotes colonization and virulence. Mol Microbiol 79:240–263.

28. Lin B, Wang Z, Malanoski AP, O’Grady EA, Wimpee CF, Vuddhakul V, Alves N, Thompson FL, Gomez-Gil B, Vora GJ. 2010. Comparative genomic analyses identify the Vibrio harveyi genome sequenced strains BAA-1116 and HY01 as Vibrio campbellii. Environ Microbiol Rep 2:81–89.

29. Urbanczyk H, Ogura Y, Hayashi T. 2013. Taxonomic revision of Harveyi clade bacteria (family Vibrionaceae) based on analysis of whole genome sequences. Int J Syst Evol Microbiol 63:2742–2751.

30. Shinoda S, Nakahara N, Uchida E, Hiraga M. 1985. Lateral Flagellar Antigen of Vibrio alginolyticus and Vibrio harveyi: Existence of Serovars Common to the Two Species. Microbiol Immunol 29:173–182.

31. Allen RD, Baumann P. 1971. Structure and arrangement of flagella in species of the genus Beneckea and Photobacterium fischeri. J Bacteriol 107:295–302.

32. Simpson CA, Podicheti R, Rusch DB, Dalia AB, van Kessel JC. 2019. Diversity in Natural Transformation Frequencies and Regulation across Vibrio Species. MBio 10:1–16.

33. Simpson CA, Petersen BD, Geyman LJ, Lee AH, Manzella MP, Papenfort K, van Kessel JC. 2020. Vibrio campbellii DS40M4 is a tractable model strain that diverges from the canonical quorum-sensing regulatory circuit in vibrios. bioRxiv 2020.03.31.019307.

34. Dalia AB, McDonough E, Camilli A. 2014. Multiplex genome editing by natural transformation. Proc Natl Acad Sci 111:8937–8942.

35. Chen M, Zhao Z, Yang J, Peng K, Baker MAB, Bai F, Lo CJ. 2017. Length-dependent flagellar growth of vibrio alginolyticus revealed by real time fluorescent imaging. Elife 6:1– 16.

36. Grossart H-P, Steward GF, Martinez J, Azam F. 2000. A Simple, Rapid Method for Demonstrating Bacterial Flagella. Appl Environ Microbiol 66:3632 LP–3636.

37. Barker CS, Meshcheryakova I V., Kostyukova AS, Samatey FA. 2010. FliO regulation of FliP in the formation of the Salmonella enterica flagellum. PLoS Genet 6.

38. Van Arnam JS, McMurry JL, Kihara M, Macnab RM. 2004. Analysis of an Engineered Salmonella Flagellar Fusion Protein, FliR-FlhB. J Bacteriol 186:2495–2498.

39. Fraser GM, Bennett JCQ, Hughes C. 1999. Substrate-specific binding of hook-associated proteins by FlgN and FliT, putative chaperones for flagellum assembly. Mol Microbiol 32:569–580.

40. Chilcott GS, Hughes KT. 2000. Coupling of Flagellar Gene Expression to Flagellar Assembly in Salmonella enterica Serovar Typhimurium andEscherichia coli. Microbiol Mol Biol Rev 64:694–708.

41. Correa NE, Peng F, Klose KE. 2005. Roles of the Regulatory Proteins FlhF and FlhG in the Vibrio cholerae Flagellar Transcription Hierarchy. J Bacteriol 187:6324LP–6332.

42. Erhardt M, Mertens ME, Fabiani FD, Hughes KT. 2014. ATPase-Independent Type-III Protein Secretion in Salmonella enterica. PLoS Genet 10.

43. Martinez RM, Jude BA, Kirn TJ, Skorupski K, Taylor RK. 2010. Role of FlgT in anchoring the flagellum of Vibrio cholerae. J Bacteriol 192:2085–2092.

44. Klose KE, Mekalanos JJ. 1998. Distinct roles of an alternative sigma factor during both free-swimming and colonizing phases of the Vibrio cholerae pathogenic cycle. Mol Microbiol 28:501–520.

45. Klose KE, Novik V, Mekalanos JJ. 1998. Identification of multiple sigma54-dependent transcriptional activators in Vibrio cholerae. J Bacteriol 180:5256–5259.

46. Kim YK, McCarter LL. 2004. Cross-regulation in Vibrio parahaemolyticus: Compensatory activation of polar flagellar genes by the lateral flagellar regulator LafK. J Bacteriol 186:4014–4018.

47. Kusumoto A, Shinohara A, Terashima H, Kojima S, Yakushi T, Homma M. 2008. Collaboration of FlhF and FlhG to regulate polar-flagella number and localization in Vibrio alginolyticus. Microbiology 154:1390–1399.

48. Correa NE, Lauriano CM, McGee R, Klose KE. 2000. Phosphorylation of the flagellar regulatory protein FlrC is necessary for Vibrio cholerae motility and enhanced colonization. Mol Microbiol 35:743–755.

49. Merino S, Shaw JG, Tomás JM. 2006. Bacterial lateral flagella: An inducible flagella system. FEMS Microbiol Lett 263:127–135.

50. Kearns DB. 2011. A field guide to bacterial swarming motility. NIH 8:634–644.

51. Dey S, Biswas M, Sen U, Dasgupta J. 2015. Unique ATPase site architecture triggers cis-mediated synchronized ATP binding in heptameric AAA+-ATPase domain of flagellar regulatory protein FlrC. J Biol Chem 290:8734–8747.

52. Chakraborty S, Biswas M, Dey S, Agarwal S, Chakrabortty T, Ghosh B, Dasgupta J. 2020. The heptameric structure of the flagellar regulatory protein FlrC is indispensable for ATPase activity and disassembled by cyclic-di-GMP. J Biol Chem 295:16960–16974.

53. Cohen EJ, Hughes KT. 2014. Rod-to-Hook transition for extracellular flagellum assembly is catalyzed by the L-ring-dependent rod scaffold removal. J Bacteriol 196:2387–2395.

54. McCarter LL. 1995. Genetic and molecular characterization of the polar flagellum of Vibrio parahaemolyticus. J Bacteriol 177:1595–1609.

55. Homma M, Fujita H, Yamaguchi S, Iino T. 1984. Excretion of unassembled flagellin by Salmonella typhimurium mutants deficient in hook-associated proteins. J Bacteriol 159:1056–1059.

56. Echazarreta MA, Kepple JL, Yen LH, Chen Y, Klose KE. 2018. A critical region in the FlaA flagellin facilitates filament formation of the Vibrio cholerae flagellum. J Bacteriol 200:1–11.

57. Kim SY, Thanh XTT, Jeong K, Kim S Bin, Pan SO, Jung CH, Hong SH, Lee SE, Rhee JH. 2014. Contribution of six flagellin genes to the flagellum biogenesis of Vibrio vulnificus and in vivo invasion. Infect Immun 82:29–42.

58. Millikan DS, Ruby EG. 2004. Vibrio fischeri Flagellin A Is Essential for Normal Motility and for Symbiotic Competence during Initial Squid Light Organ Colonization. J Bacteriol 186:4315 LP – 4325.

59. Milton DL, O’Toole R, Horstedt P, Wolf-Watz H. 1996. Flagellin A is essential for the virulence of Vibrio anguillarum. J Bacteriol 178:1310 LP – 1319.

60. Martínez-García E, Nikel PI, Chavarría M, de Lorenzo V. 2014. The metabolic cost of flagellar motion in Pseudomonas putidaKT2440. Environ Microbiol 16:291–303.

61. Ni B, Colin R, Link H, Endres RG, Sourjik V. 2020. Growth-rate dependent resource investment in bacterial motile behavior quantitatively follows potential benefit of chemotaxis. Proc Natl Acad Sci 117:595 LP – 601.

62. Ni B, Ghosh B, Paldy FS, Colin R, Heimerl T, Sourjik V. 2017. Evolutionary Remodeling of Bacterial Motility Checkpoint Control. Cell Rep 18:866–877.

63. Li W, Godzik A. 2006. Cd-hit: A fast program for clustering and comparing large sets of protein or nucleotide sequences. Bioinformatics 22:1658–1659.

64. Dalia TN, Hayes CA, Stolyar S, Marx CJ, McKinlay JB, Dalia AB. 2017. Multiplex Genome Editing by Natural Transformation (MuGENT) for Synthetic Biology in Vibrio natriegens. ACS Synth Biol 6:1650–1655.

65. Rutherford ST, van Kessel JC, Shao Y, Bassler BL. 2011. AphA and LuxR/HapR reciprocally control quorum sensing in vibrios. Genes Dev 25:397–408.

66. Chaparian RR, Olney SG, Hustmyer CM, Rowe-Magnus DA, van Kessel JC. 2016. Integration host factor and LuxR synergistically bind DNA to coactivate quorum-sensing genes in Vibrio harveyi. Mol Microbiol 101:823–840.

67. Papenfort K, Förstner KU, Cong JP, Sharma CM, Bassler BL. 2015. Differential RNA-seq of Vibrio cholerae identifies the VqmR small RNA as a regulator of biofilm formation. Proc Natl Acad Sci U S A 112:E766–E775.

