## Supplementary Info for "The polar flagellar transcriptional regulatory network in *Vibrio campbellii* deviates from canonical *Vibrio* species"

Supplemental Figures S1
Supplemental Tables S1-S3
Supplemental Datasets S1-S2

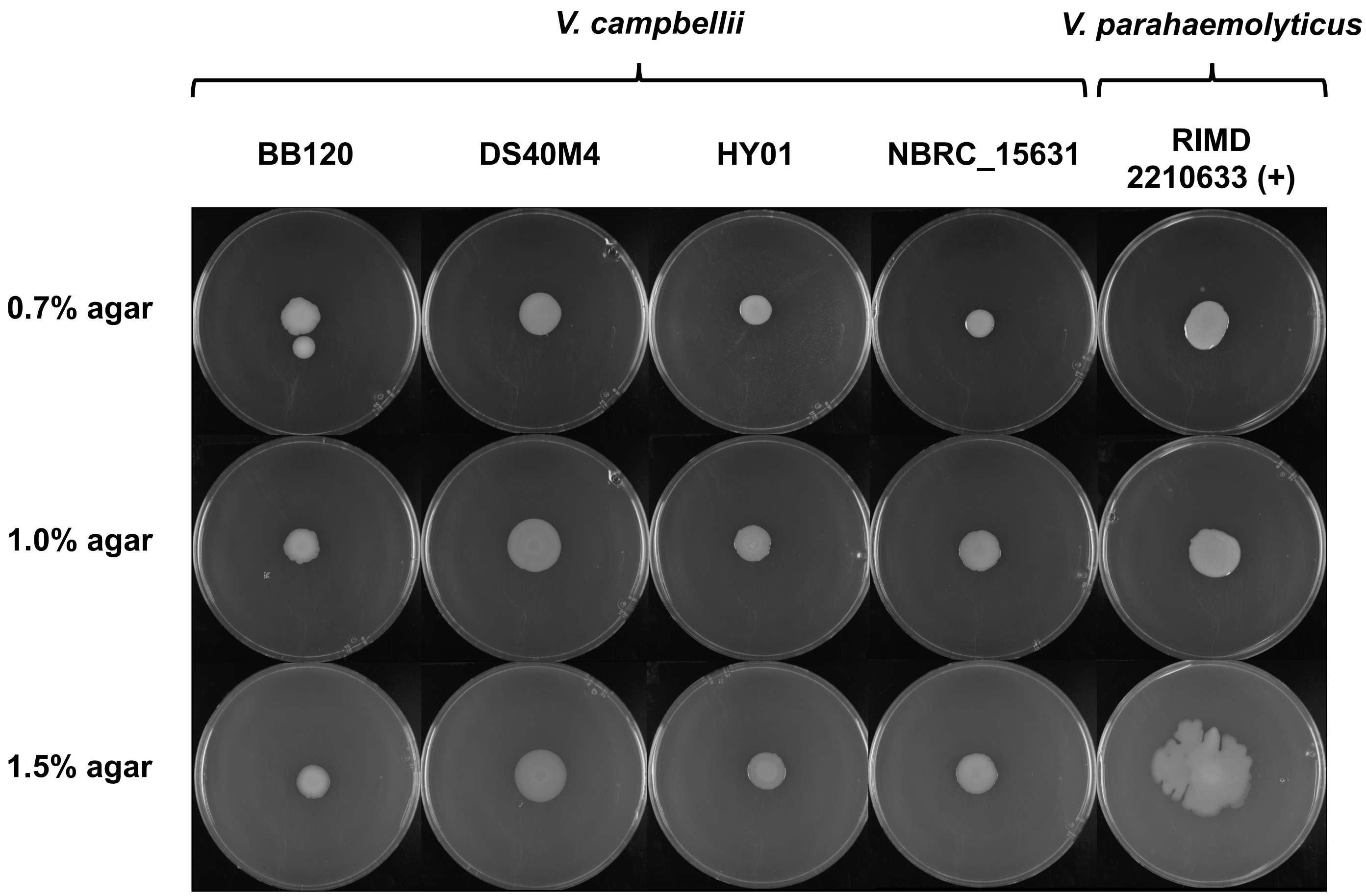

**Supplemental Fig. 1**. **Swarming motility in *V. campbellii***
Hard agar (0.7, 1.0, or 1.5%) swarm plates showing swarming phenotypes for each indicated *Vibrio* strain: *V. campbellii* BB120, DS40M4, HY01, and NBRC_15631 or *V. parahaemolyticus* RIMD 2210633. All plates were incubated at 30°C for 24h.

**Dataset S1:**
RNA-seq differential expression analysis from experiments comparing *V. campbellii* DS40M4 strains wild-type, Δ*rpoN*, Δ*flrA*, Δ*flrC*, and Δ*fliA* strains. NCBI GEO accession: GSE167483

**Dataset S2:**
Transcriptional start site analysis of data from differential RNA-seq experiments performed with *V. campbellii* DS40M4 wild-type and Δ*luxR* strains. NCBI GEO accession: GSE147616

**Table S1. Strains used in this study.**

| Name | Description | Reference |
|---|---|---|
| BB120 | *V. campbellii* type strain, derivative of BB7 (wild-type ) | (1) |
| DS40M4 | Wild-type *V. campbellii* | (2) |
| cas0034 | DS40M4, pMMB67EH-tfox-kanR | (3) |
| cas0107 | DS40M4, pMMB67EH-tfox-kanR, Δ*luxB*::*spec*$^R$ | This study |
| BDP107 | DS40M4, pMMB67EH-tfox-kanR, Δ*luxB*::*spec*$^R$, Δ*pomA* | This study |
| BDP033 | DS40M4, pMMB67EH-tfox-kanR, Δ*luxB*::*spec*$^R$, Δ*pomB* | This study |
| BDP114 | DS40M4, pMMB67EH-tfox-kanR, Δ*luxB*::*spec*$^R$, Δ*flgT* | This study |
| BDP135 | DS40M4, pMMB67EH-tfox-kanR, Δ*luxB*::*spec*$^R$, Δ*flgO* | This study |
| BDP116 | DS40M4, pMMB67EH-tfox-kanR, Δ*luxB*::*spec*$^R$, Δ*flgP* | This study |
| BDP117 | DS40M4, pMMB67EH-tfox-kanR, Δ*luxB*::*spec*$^R$, Δ*flgN* | This study |
| BDP045 | DS40M4, pMMB67EH-tfox-kanR, Δ*luxB*::*spec*$^R$, Δ*flgM* | This study |
| BDP117 | DS40M4, pMMB67EH-tfox-kanR, Δ*luxB*::*spec*$^R$, Δ*flgA* | This study |
| BDP134 | DS40M4, pMMB67EH-tfox-kanR, Δ*luxB*::*spec*$^R$, Δ*cheV* | This study |
| BDP113 | DS40M4, pMMB67EH-tfox-kanR, Δ*luxB*::*spec*$^R$, Δ*cheR* | This study |
| BDP144 | DS40M4, pMMB67EH-tfox-kanR, Δ*luxB*::*spec*$^R$, Δ*flgB* | This study |
| BDP108 | DS40M4, pMMB67EH-tfox-kanR, Δ*luxB*::*spec*$^R$, Δ*flgC* | This study |
| BDP109 | DS40M4, pMMB67EH-tfox-kanR, Δ*luxB*::*spec*$^R$, Δ*flgD* | This study |
| BDP110 | DS40M4, pMMB67EH-tfox-kanR, Δ*luxB*::*spec*$^R$, Δ*flgE* | This study |
| BDP111 | DS40M4, pMMB67EH-tfox-kanR, Δ*luxB*::*spec*$^R$, Δ*flgF* | This study |
| BDP112 | DS40M4, pMMB67EH-tfox-kanR, Δ*luxB*::*spec*$^R$, Δ*flgG* | This study |
| BDP125 | DS40M4, pMMB67EH-tfox-kanR, Δ*luxB*::*spec*$^R$, Δ*flgH* | This study |
| BDP146 | DS40M4, pMMB67EH-tfox-kanR, Δ*luxB*::*spec*$^R$, Δ*flgI* | This study |
| BDP123 | DS40M4, pMMB67EH-tfox-kanR, Δ*luxB*::*spec*$^R$, Δ*flgJ* | This study |
| BDP098 | DS40M4, pMMB67EH-tfox-kanR, Δ*luxB*::*spec*$^R$, Δ*flgK* | This study |

| | | |
|---|---|---|
| BDP105 | DS40M4, pMMB67EH-tfox-kanR, Δ*luxB*::*spec*$^{R}$, Δ*flgL* | This study |
| CAS0130 | DS40M4, pMMB67EH-tfox-kanR, Δ*luxB*::*spec*$^{R}$, Δ*flaC* | This study |
| CAS0149 | DS40M4, pMMB67EH-tfox-kanR, Δ*luxB*::*spec*$^{R}$, Δ*flaB* | This study |
| CAS0142 | DS40M4, pMMB67EH-tfox-kanR, Δ*luxB*::*spec*$^{R}$, Δ*flaE* | This study |
| BDP104 | DS40M4, pMMB67EH-tfox-kanR, Δ*luxB*::*spec*$^{R}$, ΔmotY | This study |
| BDP118 | DS40M4, pMMB67EH-tfox-kanR, Δ*luxB*::*spec*$^{R}$, Δ*flrD* | This study |
| BDP136 | DS40M4, pMMB67EH-tfox-kanR, Δ*luxB*::*spec*$^{R}$, Δ*cheW* | This study |
| BDP119 | DS40M4, pMMB67EH-tfox-kanR, Δ*luxB*::*spec*$^{R}$, Δ*ORF2* | This study |
| BDP120 | DS40M4, pMMB67EH-tfox-kanR, Δ*luxB*::*spec*$^{R}$, Δ*ORF1* | This study |
| BDP145 | DS40M4, pMMB67EH-tfox-kanR, Δ*luxB*::*spec*$^{R}$, Δ*cheB* | This study |
| BDP137 | DS40M4, pMMB67EH-tfox-kanR, Δ*luxB*::*spec*$^{R}$, Δ*cheA* | This study |
| BDP121 | DS40M4, pMMB67EH-tfox-kanR, Δ*luxB*::*spec*$^{R}$, Δ*cheZ* | This study |
| BDP101 | DS40M4, pMMB67EH-tfox-kanR, Δ*luxB*::*spec*$^{R}$, Δ*cheY* | This study |
| BDP046 | DS40M4, pMMB67EH-tfox-kanR, Δ*luxB*::*spec*$^{R}$, Δ*fliA* | This study |
| ML006 | DS40M4, pMMB67EH-tfox-kanR, Δ*luxB*::*spec*$^{R}$, Δ*flhG* | This study |
| BDP100 | DS40M4, pMMB67EH-tfox-kanR, Δ*luxB*::*spec*$^{R}$, Δ*flhF* | This study |
| BDP126 | DS40M4, pMMB67EH-tfox-kanR, Δ*luxB*::*spec*$^{R}$, Δ*flhA* | This study |
| BDP106 | DS40M4, pMMB67EH-tfox-kanR, Δ*luxB*::*spec*$^{R}$, Δ*flhB* | This study |
| BDP122 | DS40M4, pMMB67EH-tfox-kanR, Δ*luxB*::*spec*$^{R}$, Δ*fliR* | This study |
| BDP127 | DS40M4, pMMB67EH-tfox-kanR, Δ*luxB*::*spec*$^{R}$, Δ*fliQ* | This study |
| BDP138 | DS40M4, pMMB67EH-tfox-kanR, Δ*luxB*::*spec*$^{R}$, Δ*fliP* | This study |
| BDP128 | DS40M4, pMMB67EH-tfox-kanR, Δ*luxB*::*spec*$^{R}$, Δ*fliO* | This study |
| BDP129 | DS40M4, pMMB67EH-tfox-kanR, Δ*luxB*::*spec*$^{R}$, Δ*fliN* | This study |
| BDP133 | DS40M4, pMMB67EH-tfox-kanR, Δ*luxB*::*spec*$^{R}$, Δ*fliM* | This study |
| BDP141 | DS40M4, pMMB67EH-tfox-kanR, Δ*luxB*::*spec*$^{R}$, Δ*fliL* | This study |

| | | |
|---|---|---|
| BDP124 | DS40M4, pMMB67EH-tfox-kanR, Δ*luxB*::*spec*$^{R}$, Δ*fliK* | This study |
| BDP142 | DS40M4, pMMB67EH-tfox-kanR, Δ*luxB*::*spec*$^{R}$, Δ*fliJ* | This study |
| BDP130 | DS40M4, pMMB67EH-tfox-kanR, Δ*luxB*::*spec*$^{R}$, Δ*fliI* | This study |
| BDP131 | DS40M4, pMMB67EH-tfox-kanR, Δ*luxB*::*spec*$^{R}$, Δ*fliH* | This study |
| BDP132 | DS40M4, pMMB67EH-tfox-kanR, Δ*luxB*::*spec*$^{R}$, Δ*fliG* | This study |
| BDP143 | DS40M4, pMMB67EH-tfox-kanR, Δ*luxB*::*spec*$^{R}$, Δ*fliF* | This study |
| BDP102 | DS40M4, pMMB67EH-tfox-kanR, Δ*luxB*::*spec*$^{R}$, Δ*fliE* | This study |
| BDP031 | DS40M4, pMMB67EH-tfox-kanR, Δ*luxB*::*spec*$^{R}$, Δ*flrC* | This study |
| BDP099 | DS40M4, pMMB67EH-tfox-kanR, Δ*luxB*::*spec*$^{R}$, Δ*flrB* | This study |
| BDP030 | DS40M4, pMMB67EH-tfox-kanR, Δ*luxB*::*spec*$^{R}$, Δ*flrA* | This study |
| BDP150 | DS40M4, pMMB67EH-tfox-kanR, Δ*luxB*::*spec*$^{R}$, Δ*fliS* | This study |
| BDP151 | DS40M4, pMMB67EH-tfox-kanR, Δ*luxB*::*spec*$^{R}$, Δ*fliS* | This study |
| BDP139 | DS40M4, pMMB67EH-tfox-kanR, Δ*luxB*::*spec*$^{R}$, Δ*fliT* | This study |
| BDP140 | DS40M4, pMMB67EH-tfox-kanR, Δ*luxB*::*spec*$^{R}$, Δ*flaG* | This study |
| CAS0128 | DS40M4, pMMB67EH-tfox-kanR, Δ*luxB*::*spec*$^{R}$, Δ*flaA* | This study |
| CAS0150 | DS40M4, pMMB67EH-tfox-kanR, Δ*luxB*::*spec*$^{R}$, Δ*flaD* | This study |
| CAS0131 | DS40M4, pMMB67EH-tfox-kanR, Δ*luxB*::*spec*$^{R}$, Δ*flaF* | This study |
| BDP029 | DS40M4, pMMB67EH-tfox-kanR, Δ*luxB*::*spec*$^{R}$, ΔrpoN | This study |
| BDP103 | DS40M4, pMMB67EH-tfox-kanR, Δ*luxB*::*spec*$^{R}$, Δ*motX* | This study |
| ML05 | DS40M4, pMMB67EH-tfox-kanR, Δ*luxB*::*spec*$^{R}$, Δ*flrA, ΔflrC* | This study |
| BDP152 | DS40M4, pMMB67EH-tfox-kanR, ΔflrA, ΔflrC, Δ*luxB*::Ptac-flrA-*Tm*$^{R}$, | This study |
| BDP153 | DS40M4, pMMB67EH-tfox-kanR, ΔflrA, ΔflrC, Δ*luxB*::Ptac-flrC-*Tm*$^{R}$, | This study |
| BDP037 | DS40M4, pMMB67EH-tfox-kanR, ΔrpoN, Δ*luxB*::rpoN-*Tm*$^{R}$, | This study |

| | | |
|---|---|---|
| BDP038 | DS40M4, pMMB67EH-tfox-kanR, ΔflrA, Δ*luxB*::flrA-*Tm*$^{R}$, | This study |
| BDP054 | DS40M4, pMMB67EH-tfox-kanR, ΔflrC, Δ*luxB*::Ptac-flrC-Spec$^{R}$, | This study |
| BDP055 | DS40M4, pMMB67EH-tfox-kanR, ΔfliA, Δ*luxB*::Ptac-fliA-SpecR, | This study |
| AY05 | DS40M4, pMMB67EH-tfox-kanR, ΔluxB::specR, ΔlafK | This study |
| AY11 | DS40M4, pMMB67EH-tfox-kanR, ΔluxB::specR, ΔflrA, ΔlafK | This study |

**Table S2. Plasmids used in this study.**

| Name | Description | Reference |
|---|---|---|
| pMMB67EH-tfoX-kanR | *kan*$^{R}$ , P*tac-tfoX*, *lacI*, IncQ origin | (3) |
| pJV298 | *CM*$^{R}$, P*tac-gfp*, *lacI*, p15a origin | (4) |

**Table S3. Oligonucleotides used in this study.**

| Name | Sequence | Notes |
|---|---|---|
| **Oligos for MuGENT or SOE products** | | |
| CAS0148 | CGTGCTCAAGTCTTCACTGATGATG | DS40M4 Δ*luxB* F1 |
| CAS0149 | GTCGACGGATCCCCGGAATGATGACTTGATCAGAAGAACGCTTTGA | DS40M4 Δ*luxB* R1 |
| CAS0150 | GAAGCAGCTCCAGCCTACACACTCGTAACGTTTAAACGATGCTGAG | DS40M4 Δ*luxB* F2 |
| CAS0151 | GGTGAATGGCCACAAGGTACCT | DS40M4 Δ*luxB* R2 |
| BP145 | GCGGCATGGATATCAATTCCATGGTC | DS40M4 delta FlrA F1 |
| BP180 | gctaattcagtttaagcggccatAAGCAGCTTTGCCAAACCTTGCATAAG | DS40M4 delta FlrA R1, universal overlap homology |
| BP182 | atggccgcttaaactgaattagcTAGGGTCACCCTGGCGTTAAAGC | DS40M4 delta FlrA F2, universal overlap homology |
| BP148 | AGAAAGGGAAACATCGGCATCACC | DS40M4 delta FlrA R2 |
| BP149 | GGTATCTAGTAAGATCACACCTGC | detect delta FlrA DS40M4, R |
| BP150 | CGCAGTATCAACCGCCTTGAAGG | DS40M4 delta RpoN F1 |
| BP151 | gctaattcagtttaagcggccatCATTCAGTGTTACTTACCTTGTAATGCTCAG | DS40M4 delta RpoN R1, universal overlap homology |

| | | |
|---|---|---|
| BP152 | atggccgcttaaactgaattagcTAGGCACCAATTGAAAAGGAAAGTCTATGC | DS40M4 delta RpoN F2 , universal overlap homology |
| BP153 | GGAGACAGAGGCAGCGGCTA | DS40M4 delta RpoN R2 |
| BP154 | GAATCAATTGCTGCATACATACTTTCG | detect delta RpoN DS40M4, R |
| BP155 | ACCTTACTTAATGAGTAGGGTGTTGTGCC | DS40M4 delta FlrC F1 |
| BP156 | gctaattcagtttaagcggccatTTGCGCCATTATGATTCTCCAGTAATTGTTATC | DS40M4 delta FlrC R1, universal overlap homology |
| BP157 | atggccgcttaaactgaattagcCCGGCTTAGTATTCACATAAATAGCCATAGG | DS40M4 delta FlrC F2 , universal overlap homology |
| BP158 | AGCAATTTCTGAATACCTTGGTCGTCC | DS40M4 delta FlrC R2 |
| BP159 | TGATGGCTTTCGTTAGGAGATCA | detect delta FlrC DS40M4, R |
| BP160 | GAAGTACAGAACCTATTGGAAATGCTCGG | DS40M4 delta FliAp F1 |
| BP161 | gctaattcagtttaagcggccatCACCAAAAGGGTCCTCTGTGAATTCAG | DS40M4 delta FliAp R1, universal overlap homology |
| BP263 | atggccgcttaaactgaattagcAGCGAATCTCGCGTAAGCCAAATACTG | DS40M4 delta FliAp F2 , universal overlap homology |
| BP163 | CCATACCTACGCCACGACCAGA | DS40M4 delta FliAp R2 |
| BP164 | CACGGATGTGTTTAAGCAGGT | detect delta FliAp DS40M4, R |
| BP234 | gtcgacggatccccggaatCTAAAGTAGGCGTTTACGCTGACTCG | DS40M4 luxB::RpoN_TmR complement R, homology to TmR cassette |
| BP235 | tcaaagcgttcttctgatcaagtcatcCTATCTGTAGAAGACAACATCATGGCCGTT | DS40M4 luxB::RpoN_TmR complement F, 500 bp upstream included homology to luxB UP, v2 |
| BP241 | tcaaagcgttcttctgatcaagtcatcGCGTAGCTGTCTATCTATGGAAGATGGTG | DS40M4 luxB::FlrA_TmR complement F, 500 bp upstream included, homology to luxB UP |
| BP242 | gtcgacggatccccggaatCTAGCGTTGAAGGTTGTACTTACGCAT | DS40M4 luxB::FlrA_TmR complement R, homology to TmR cassette |
| BP256 | aattcggcgtaagcttaaggagatatacatATGGCGCAAAGCAAAGTGCTG | DS40M4 FlrC Complement F, |

| | | |
|---|---|---|
| | | homology to upstream+Ptac from pJV298 |
| BP244 | gtcgacggatccccggaatCTAAGCCGGAATATCGATACCAGCGTC | DS40M4 luxB::FlrC_TmR complement R, homology to TmR cassette |
| BP257 | attcggcgtaagcttaaggagatatacatGTGAATAAAGCGATTACCTATGACCAACATG | DS40M4 FliAp Complement F, homology to upstream+Ptac from pJV298 |
| BP246 | gtcgacggatccccggaatTTAGTCATTTTGTGTCCACGCACTGAG | DS40M4 luxB::FliAp_TmR complement R, homology to TmR cassette |
| BP247 | cgcctggagagtgtatagcgattac | detect delta luxB DS40M4, F |
| BP248 | catctcagttgcctccttcattattagctg | detect delta luxB DS40M4, R |
| BP264 | GCCGTTTTGCCGCTTCAGTTAC | DS40M4 delta FlgM F1 |
| BP265 | gtcgacggatccccggaatTATACCTGCCATATTTCAACCTTAAACTCG | DS40M4 delta FlgM R1, Ab overlap homology |
| BP266 | gaagcagctccagcctacaTAACTGACTTAGGCCCAAGGTAAGC | DS40M4 delta FlgM F2 , Ab overlap homology |
| BP267 | TGTTCATTACGTCGCCAATCTGC | DS40M4 delta FlgM R2 |
| BP268 | CCTGACTTAGGCTTTCGTCTTCGG | detect delta FlgM DS40M4, R |
| BP374 | CACTTGCTCATTCAAATAAGGCTGAATCG | DS40M4 delta fliK F1 |
| BP375 | gctaattcagtttaagcggccatGTGGTTTCAAAGCGTGATGGAATTAG | DS40M4 delta fliK R1, universal overlap homology |
| BP376 | atggccgcttaaactgaattagcCATATAGAAACTCTCGTCATCGGCAGC | DS40M4 delta fliK F2, universal overlap homology |
| BP377 | CTGACAGAAGAACAGATTGAGTTGATCAAG | DS40M4 delta fliK R2 |
| BP378 | ATTGATTGAGCAAGGTCTCTGTTTACTC | DS40M4 delta fliK detect, R, 250 bp downstream |
| BP379 | CAGCGCTGCGGTTCGATCATC | DS40M4 delta fliE F1 |
| BP380 | gctaattcagtttaagcggccatGTAATTTTAGGTAATAAGTGTGGCAGACAAGTC | DS40M4 delta fliE R1, universal overlap homology |
| BP381 | atggccgcttaaactgaattagcTGTCACCTCTAAGCCAAAAGTTTGAC | DS40M4 delta fliE F2 , universal overlap homology |
| BP382 | CGATCTGGACCACAACATGAATGC | DS40M4 delta fliE R2 |

| | | |
|---|---|---|
| BP383 | TGGAAACGCTGATCGAATGCAACGG | detect delta fliE DS40M4, R, 250 bp downstream |
| BP384 | TTTTGAACGATTCATCAGCAACGCC | DS40M4 delta flhA F1 |
| BP385 | gctaattcagtttaagcggccatTAACCCGCGCATCATTAATGGATTTATAG | DS40M4 delta flhA R1, universal overlap homology |
| BP386 | atggccgcttaaactgaattagcCATACTGCGAGGTGTCTCAATCGG | DS40M4 delta flhA F2 , universal overlap homology |
| BP387 | CGTGGTGGTGATGTCCATCTTGCG | DS40M4 delta flhA R2 |
| BP388 | CGTCACTAATGAAAATAATTGCATGGTGTG | detect delta flhA DS40M4, R, 250 bp downstream |
| BP389 | AACAAGCTGCTCAGAAAGCAGCTC | DS40M4 delta flhF F1 |
| BP390 | gctaattcagtttaagcggccatGGGCGGCGGACTACTATGACTG | DS40M4 delta flhF R1, universal overlap homology |
| BP391 | atggccgcttaaactgaattagcCAAACTATAAATCCATTAATGATGCGCGGG | DS40M4 delta flhF F2 , universal overlap homology |
| BP392 | GCGTCGCAAGCGGATTTGATTCAG | DS40M4 delta flhF R2 |
| BP393 | CCGGTATTGAACCGGGATTGGCTG | detect delta flhF DS40M4, R, 250 bp downstream |
| BP394 | ATCACGACCATGTCTGGTTGTGCG | DS4 delta flgK F1 |
| BP395 | gctaattcagtttaagcggccatATCTGACGCCATACATGCCCC | DS4 delta flgK R1, universal overlap homology |
| BP396 | atggccgcttaaactgaattagcAGGAGGCCTTAGATGTCAACGCG | DS4 delta flgK F2, universal overlap homology |
| BP397 | GGCTAGGTTGGTAGCCAGAGCC | DS4 delta flgK R2 |
| BP398 | ACTTCTTGGTTTTCTAGACGGTTACGAAC | DS4 delta flgK detect, R, 250bp downstream |
| BP399 | ACTGAGCCTTGACCAGTCAATGTGC | DS4 delta flrB (flaL) F1 |
| BP400 | gctaattcagtttaagcggccatTAAGCGGCTTTTGTCGATAACAATTACTG | DS4 delta flrB (flaL) R1, universal overlap homology |
| BP401 | atggccgcttaaactgaattagcCATTCACTGCCTGTCGAAATATCGC | DS4 delta flrB (flaL) F2, universal overlap homology |
| BP402 | TTAGAAATCAACTATGACATGTTGGATCGC | DS4 delta flrB (flaL) R2 |
| BP403 | TAGGGATGCGACGCACAACACTG | DS4 delta flrB (flaL) detect, R, 250bp downstream |
| BP404 | TAAGTGAACACAAACCGTTTGATAGCAC | DS4 delta motX F1 |
| BP405 | gctaattcagtttaagcggccatCGCGACACCTTCTGGTAGATGTACTC | DS4 delta motX R1, universal overlap homology |

| | | |
|---|---|---|
| BP406 | atggccgcttaaactgaattagcGTGTAGATGGGGACAGGCGC | DS4 delta motX F2, universal overlap homology |
| BP407 | CGATTCGGTCGTGTCTTAAAAGACAATAAC | DS4 delta motX R2 |
| BP408 | AAAATGCGACTCACTGTGGCAAATATTTG | DS4 delta motX detect, R, 250bp downstream |
| BP409 | CTATGGTATCTGCGCCCGACCTAGG | DS4 delta motY F1 |
| BP410 | gctaattcagtttaagcggccatCGCACCCAGGTTTGATCTTAAATATTCAC | DS4 delta motY R1, universal overlap homology |
| BP411 | atggccgcttaaactgaattagcCATCTAGGTACTTCTCTCAAACGGCAG | DS4 delta motY F2, universal overlap homology |
| BP412 | ACATGATCCAATGTGCAGCTGGTG | DS4 delta motY R2 |
| BP413 | AAACCGCAATTTGGGAGAAATGTAAAATTC | DS4 delta motY detect, R, 250bp downstream |
| BP414 | ATATCGGTTTTATTCATGACATCAGTTCCC | DS4 delta flgL F1 |
| BP415 | gctaattcagtttaagcggccatCATCTAAGGCCTCCTTTAGCGCAAC | DS4 delta flgL R1, universal overlap homology |
| BP416 | atggccgcttaaactgaattagcTAACGCGCCTATAGTCTATGCAGTGC | DS4 delta flgL F2, universal overlap homology |
| BP417 | TGATAGCCACAGATTCTTGTGCGCC | DS4 delta flgL R2 |
| BP418 | ATAAACAACAAGTTATAGCGCATATGGGG | DS4 delta flgL detect, R, 250bp downstream |
| BP419 | ATTGTCTAATGGCTTGATCACCACTTC | DS4 delta cheY (DSB67_11290) F1 |
| BP420 | gctaattcagtttaagcggccatTTTACGGCTGCAACGCTAAAAGAAAAAC | DS4 delta cheY DSB67_11290) R1, universal overlap homology |
| BP421 | atggccgcttaaactgaattagcGCGCATTGTTGAGAAATCATCAACAATAAGG | DS4 delta cheY (DSB67_11290) F2, universal overlap homology, v2 |
| BP422 | GGCGCAGCTCAACCAAATCG | DS4 delta cheY (DSB67_11290) R2, v2 |
| BP423 | AAGCACTGGTTGAATCAATAAAACAACTTC | DS4 delta cheY (DSB67_11290) detect, R, 250bp downstream |
| BP424 | GAATCGACGTCATCTCTTCGCGCATG | DS4 delta flhB F1 |
| BP425 | gctaattcagtttaagcggccatCGTCACTAATGAAAATAATTGCATGGTGTG | DS4 delta flhB R1, universal overlap homology |
| BP426 | atggccgcttaaactgaattagcCAATCCAGCCTCCTAGCAATCCAAG | DS4 delta flhB F2, universal overlap homology |
| BP427 | AGATAAAATGGCGGTACAAATCTCCG | DS4 delta flhB R2 |

| | | |
|---|---|---|
| BP428 | TTAAGACGGCACTCAGTATGTCGTTG | DS4 delta flhB detect, R, 250bp downstream |
| BP429 | AGCTGCTTCTAGCGCGTTACCTAAG | DS4 delta pomA F1 |
| BP430 | gctaattcagtttaagcggccatACAAAGCACTCCTTTCGCTATACTTAAAC | DS4 delta pomA R1, universal overlap homology |
| BP431 | atggccgcttaaactgaattagcTCCGGAGGTCATCTGATGGATGAAG | DS4 delta pomA F2, universal overlap homology |
| BP432 | CGTTGTTGCGATGTTGTGTTTGC | DS4 delta pomA R2 |
| BP433 | AACTCTTGCGCAATGATGCTGGTG | DS4 delta pomA detect, R, 250bp downstream |
| BP434 | ATCAAGAACTCAGATCCAGCGTACG | DS4 delta flgT F1 |
| BP435 | gctaattcagtttaagcggccatGCAACACGTATAATGCCTTGCGATATG | DS4 delta flgT R1, universal overlap homology |
| BP436 | atggccgcttaaactgaattagcTGTTAGTAACCGTTCGGGTATAAATATTGC | DS4 delta flgT F2, universal overlap homology |
| BP437 | ACAAAGTACGCTAAGTCTATCGGCATTTTG | DS4 delta flgT R2 |
| BP438 | CTTTACCATTGTAAATTGGTGAGTAGGCG | DS4 delta flgT detect, R, 250bp downstream |
| BP439 | GACAACCTAGGTGACTTTGACCAAAC | DS4 delta flgO F1 |
| BP440 | gctaattcagtttaagcggccatGCCATCTTTTCATGGTTTGATTCTCCAG | DS4 delta flgO R1, universal overlap homology |
| BP441 | atggccgcttaaactgaattagcGCCCGTAGGAAACTAGCATGAAGAAG | DS4 delta flgO F2, universal overlap homology |
| BP442 | CTTCTTGTGCCATTTCAAGCAGC | DS4 delta flgO R2 |
| BP443 | TGATCGGCAAGTTCTGCACGCC | DS4 delta flgO detect, R, 250bp downstream |
| BP444 | GGTGAAAACAACCCAATCGCTTCAAAC | DS4 delta flgP F1 |
| BP445 | gctaattcagtttaagcggccatTGCTAGTTTCCTACGGGCGGAGAATC | DS4 delta flgP R1, universal overlap homology |
| BP446 | atggccgcttaaactgaattagcAGCAGACCTTGTTCTAATACAACTTCAC | DS4 delta flgP F2, universal overlap homology |
| BP447 | TTGCGGCCAGTCACGACATCAC | DS4 delta flgP R2 |
| BP448 | GGTAAGGTCGGCATGACCTACAATG | DS4 delta flgP detect, R, 250bp downstream |
| BP449 | AACCGCGACACCTGCTATATAGC | DS4 delta flgN F1 |
| BP450 | gctaattcagtttaagcggccatAACTCTTTGAATCAATTCCACGACTACTC | DS4 delta flgN R1, universal overlap homology |
| BP451 | atggccgcttaaactgaattagcGAATGCTTACCTTGGGCCTAAGTC | DS4 delta flgN F2, universal overlap homology |

| | | |
|---|---|---|
| BP452 | CAAATTTGTCCGTTAAACCCGTCAGTG | DS4 delta flgN R2 |
| BP453 | AAAAGCGCCAGCACAACAAGACG | DS4 delta flgN detect, R, 250bp downstream |
| BP454 | GTTTTGCCGTCAAAGACCTTCATCTC | DS4 delta flgA F1 |
| BP455 | gctaattcagtttaagcggccatGGGTTGGCTGTTGTACGGCAAATC | DS4 delta flgA R1, universal overlap homology |
| BP456 | atggccgcttaaactgaattagcACTGTTTCTCTGGTATTATGCCTGCC | DS4 delta flgA F2, universal overlap homology |
| BP457 | GCAGCAGCATTTGTTTACTAGAATCCG | DS4 delta flgA R2 |
| BP458 | CTAGTGGATGCAGATTTGGCATCG | DS4 delta flgA detect, R, 250bp downstream |
| BP459 | GAAAGAGCAGTCACACCTAAGATTATCTG | DS4 delta cheV F1 |
| BP460 | gctaattcagtttaagcggccatCGTCATATGCTCATCTCCATCTCAAAC | DS4 delta cheV R1, universal overlap homology |
| BP461 | atggccgcttaaactgaattagcGGTAACGCGGTTAAATCTGCATTAACAC | DS4 delta cheV F2, universal overlap homology |
| BP462 | TGTAAATGCTTGAACCTTCGTGGAAC | DS4 delta cheV R2 |
| BP463 | CCACAATGTTTCGTTAGTCGTCATCG | DS4 delta cheV detect, R, 250bp downstream |
| BP464 | GGCATTGAAAGTACATCGGGCGC | DS4 delta cheR F1 |
| BP465 | gctaattcagtttaagcggccatAAAGTCACGATACTCTTGATCACTGATTG | DS4 delta cheR R1, universal overlap homology |
| BP466 | atggccgcttaaactgaattagcCAACACGCTTACTAGCTGACTCTTTTAATG | DS4 delta cheR F2, universal overlap homology |
| BP467 | AACACGCACTAGCGCCACACG | DS4 delta cheR R2 |
| BP468 | AAGCCAAAGTGATTGATGGAGTGTC | DS4 delta cheR detect, R, 250bp downstream |
| BP469 | TTTTATTGATCGATTTGCCGTACAACAGC | DS4 delta flgB F1 |
| BP470 | gctaattcagtttaagcggccatTGTTTGCCTCTACAGTAAGAACTGACC | DS4 delta flgB R1, universal overlap homology |
| BP471 | atggccgcttaaactgaattagcAGCAATCAAAGGGGAATAATTTAGATGAGC | DS4 delta flgB F2, universal overlap homology |
| BP472 | AGTTATGTCCTGGACGTTCCGTCATG | DS4 delta flgB R2 |
| BP473 | GTTGTACTCCGCATTCAGCGGTTTG | DS4 delta flgB detect, R, 250bp downstream |
| BP474 | ACAATCATTGGTACAGACGCTGAGATC | DS4 delta flgC F1 |
| BP475 | gctaattcagtttaagcggccatAGCTCATCTAAATTATTCCCCTTTGATTGC | DS4 delta flgC R1, universal overlap homology |
| BP476 | atggccgcttaaactgaattagcGCAGATGGGTCAATAAGGATAAGGGG | DS4 delta flgC F2, universal overlap homology |

| | | |
|---|---|---|
| BP477 | TTCAATCGCACCGGTTTGGATGC | DS4 delta flgC R2 |
| BP478 | CTTGCCGATGCCATCTACGGTCG | DS4 delta flgC detect, R, 250bp downstream |
| BP479 | AAAAGTTTGAGATGGAGATGAGCATATGAC | DS4 delta flgD F1 |
| BP480 | gctaattcagtttaagcggccatTTCCGGCCATACGCTACCCC | DS4 delta flgD R1, universal overlap homology |
| BP481 | atggccgcttaaactgaattagcTTCAATAGCGCGTCAGGTTAGGAG | DS4 delta flgD F2, universal overlap homology |
| BP482 | TTCTGACCGATTGGCTCAAGACCAC | DS4 delta flgD R2 |
| BP483 | AATGCTTGAACCTTCGTGGAACTG | DS4 delta flgD detect, R, 250bp downstream |
| BP484 | TGGTCATTTCAGATATCGAAATGCCAG | DS4 delta flgE F1 |
| BP485 | gctaattcagtttaagcggccatTTCCAAAATCTCCTAACCTGACGCG | DS4 delta flgE R1, universal overlap homology |
| BP486 | atggccgcttaaactgaattagcAGCGCTAGTTAGATAGCTCTATCATCTG | DS4 delta flgE F2, universal overlap homology |
| BP487 | GGCAATGTTGCCGTTACCATAACTG | DS4 delta flgE R2 |
| BP488 | TAAATTGAGGGTTTTCAGATGGCAATTCAG | DS4 delta flgE detect, R, 250bp downstream |
| BP489 | TTCCTGCCACTACAACCTTGGTCG | DS4 delta flgF F1 |
| BP490 | gctaattcagtttaagcggccatCGCGATCCATAAATTACTCCAAAAGTTCTC | DS4 delta flgF R1, universal overlap homology |
| BP491 | atggccgcttaaactgaattagcCAGAGTTAGAGGTTCACAATGCATCC | DS4 delta flgF F2, universal overlap homology |
| BP492 | TGACGGTAGAACTTTTCGTTTTGGC | DS4 delta flgF R2 |
| BP493 | TGCGTCGCAACCACTTTAGAACCG | DS4 delta flgF detect, R, 250bp downstream |
| BP494 | AAGCAATTGTCTCAGCAGGACCC | DS4 delta flgG F1 |
| BP495 | gctaattcagtttaagcggccatGCATTGTGAACCTCTAACTCTGTTAACTC | DS4 delta flgG R1, universal overlap homology |
| BP496 | atggccgcttaaactgaattagcTTACTAAGATCGGGATTTAGGGAGCC | DS4 delta flgG F2, universal overlap homology |
| BP497 | TAGCAAAAGGTGTGCCAGTTTAGG | DS4 delta flgG R2 |
| BP498 | TTTTGTTTAGGATGAATCGGTGCCC | DS4 delta flgG detect, R, 250bp downstream |
| BP499 | TTTACACCAACAACCCAATGGATCTG | DS4 delta flgH F1 |
| BP500 | gctaattcagtttaagcggccatTCATTGGCTCCCTAAATCCCGATC | DS4 delta flgH R1, universal overlap homology |
| BP501 | atggccgcttaaactgaattagcGACGTCGAAAAGACGGCAAATTGAC | DS4 delta flgH F2, universal overlap homology |

| | | |
|---|---|---|
| BP502 | TGAACATCTGGTGAGCCATCAATCATC | DS4 delta flgH R2 |
| BP503 | AATGCCGAAGTTTTGCAGCATTGCC | DS4 delta flgH detect, R, 250bp downstream |
| BP504 | GTGTGGCGCTAGTGCGTGTTC | DS4 delta flgI F1 |
| BP505 | gctaattcagtttaagcggccatTGCTTACTCAATCTCTTGTAGTCAATTTGC | DS4 delta flgI R1, universal overlap homology |
| BP506 | atggccgcttaaactgaattagcTCGATTAGGAGAGAAACCATGGTCAAG | DS4 delta flgI F2, universal overlap homology |
| BP507 | TGAAGGTGTCATTGGCTGCCTGC | DS4 delta flgI R2 |
| BP508 | ATTTGACGGTAGAACTTTTCGTTTTGGC | DS4 delta flgI detect, R, 250bp downstream |
| BP509 | TTTAAAGATACCAATGGCTTATTCCGCC | DS4 delta flgJ F1 |
| BP510 | gctaattcagtttaagcggccatACCATGGTTTCTCTCCTAATCGATTAGATG | DS4 delta flgJ R1, universal overlap homology |
| BP511 | atggccgcttaaactgaattagcTTCACCCGATTACGAATGCTTGATAGAC | DS4 delta flgJ F2, universal overlap homology |
| BP512 | ATGCGTCAGATACTGAGCTCTCTG | DS4 delta flgJ R2 |
| BP513 | AGAAATGTTATGACCAGTCGTATTCAACTG | DS4 delta flgJ detect, R, 250bp downstream |
| BP514 | CTCATGAGTCTTCTTCATTGCTTCTGC | DS4 delta ORF3 (flrD) F1 |
| BP515 | gctaattcagtttaagcggccatCGCGGAAGACGTCGATAAAGATTG | DS4 delta ORF3 (flrD) R1, universal overlap homology |
| BP516 | atggccgcttaaactgaattagcGATTCATCAGCCAATTACAGGTGAGC | DS4 delta ORF3 (flrD) F2, universal overlap homology |
| BP517 | GTCCGATGGCAGCGAAATTCCG | DS4 delta ORF3 (flrD) R2 |
| BP518 | AACACGCGTATCATCGTGATTGAG | DS4 delta ORF3 (flrD) detect, R, 250bp downstream |
| BP519 | TTTCAACAATGCATCCCCACGACGG | DS4 delta cheW F1 |
| BP520 | gctaattcagtttaagcggccatATTGGCTGATGAATCATGGCTGAAG | DS4 delta cheW R1, universal overlap homology |
| BP521 | atggccgcttaaactgaattagcAGTTAATCCTCGTTAATGTCGTTCCCG | DS4 delta cheW F2, universal overlap homology |
| BP522 | AGATGGCGATTAAAGTACTAGTCGTTGATG | DS4 delta cheW R2 |
| BP523 | ACGTCGTCATGCTTGGTGAAAGCATG | DS4 delta cheW detect, R, 250bp downstream |
| BP524 | GTGCCCCACATCGGTTTGCC | DS4 delta ORF2 F1 |

| | | |
|---|---|---|
| BP525 | gctaattcagtttaagcggccatTGCTAAACGCAGGTTTAGATGTAAAATCAC | DS4 delta ORF2 R1, universal overlap homology |
| BP526 | atggccgcttaaactgaattagcTTATTGCTCATCGAACGCTAACCTCTC | DS4 delta ORF2 F2, universal overlap homology |
| BP527 | GACGAAGAAATGTCGAAAGCGGTAG | DS4 delta ORF2 R2 |
| BP528 | GTGGCTTCAAATTGACTATCGTTCCG | DS4 delta ORF2 detect, R, 250bp downstream |
| BP529 | GTTCAAGATATCCGCTTGACGGCG | DS4 delta ORF1 F1 |
| BP530 | gctaattcagtttaagcggccatGATGAGCAATAACGACATACTTTCCAGTG | DS4 delta ORF1 R1, universal overlap homology |
| BP531 | atggccgcttaaactgaattagcGATCTTTCCTACGCTAAGCCCACTTC | DS4 delta ORF1 F2, universal overlap homology |
| BP532 | GAAGTGGTCGCTACAGTTGAAGCGC | DS4 delta ORF1 R2 |
| BP533 | GGTTCCGCTGCAAAAGTCTACGG | DS4 delta ORF1 detect, R, 250bp downstream |
| BP534 | TTTTCAATCTTTATCGACGTCTTCCGC | DS4 delta cheB F1 |
| BP535 | gctaattcagtttaagcggccatGGAAAGATCATGATCGTTTGGAGTGTAG | DS4 delta cheB R1, universal overlap homology |
| BP536 | atggccgcttaaactgaattagcCTTATTCCTTTTATCGACTCCAGCTTCG | DS4 delta cheB F2, universal overlap homology |
| BP537 | CGCTCGTTCAGATGCTAGAAAGTGG | DS4 delta cheB R2 |
| BP538 | GGCCACGGCCATGTAGTTATCGTAC | DS4 delta cheB detect, R, 250bp downstream |
| BP539 | AGTGATTTTACATCTAAACCTGCGTTTAGC | DS4 delta cheA F1 |
| BP540 | gctaattcagtttaagcggccatAGCGAAGCTGGAGTCGATAAAAGG | DS4 delta cheA R1, universal overlap homology |
| BP541 | atggccgcttaaactgaattagcACGTTACCTTTAAAATCCAAGACTGGAC | DS4 delta cheA F2, universal overlap homology |
| BP542 | AATACATGATTGCGAAAGCAAATGAACTG | DS4 delta cheA R2 |
| BP543 | GTCAAATCATTCGTCGCGTGATCAC | DS4 delta cheA detect, R, 250bp downstream |
| BP544 | CACTTGGATTTTGCAAAGCGTGTTG | DS4 delta cheZ F1 |
| BP545 | gctaattcagtttaagcggccatAAGGTAACGTATGAGCTACGATTTAGACG | DS4 delta cheZ R1, universal overlap homology |
| BP546 | atggccgcttaaactgaattagcCTCTAGTGAGATCATCCTGAATAGACCC | DS4 delta cheZ F2, universal overlap homology |
| BP547 | AAGAGGTAGAGCGTCGTGAGCC | DS4 delta cheZ R2 |

| | | |
|---|---|---|
| BP548 | ATGATCACAGCGGAAGCAAAGCG | DS4 delta cheZ detect, R, 250bp downstream |
| BP549 | CCACACCAAGCTGAAGTTGGTCAGG | DS4 delta fliR F1 |
| BP550 | gctaattcagtttaagcggccatGGAGGCTGGATTGGCAGAGTCAG | DS4 delta fliR R1, universal overlap homology |
| BP551 | atggccgcttaaactgaattagcACGCGCCTCTAGTAAAGCACCTG | DS4 delta fliR F2, universal overlap homology |
| BP552 | GATTGTCCGTGGTCGGATGCCG | DS4 delta fliR R2 |
| BP553 | TTCCGTGACGCACTGTGGATGG | DS4 delta fliR detect, R, 250bp downstream |
| BP554 | ATGTCTTCATCAGTATTTTGACGCGTTAC | DS4 delta fliQ F1 |
| BP555 | gctaattcagtttaagcggccatGAGGCGCGTATGGAGTATCCGAC | DS4 delta fliQ R1, universal overlap homology |
| BP556 | atggccgcttaaactgaattagcGTCTTACCCTCTTCTCTCTGTGCC | DS4 delta fliQ F2, universal overlap homology |
| BP557 | CGGTAGAGCAACTGCGTTCACCAAC | DS4 delta fliQ R2 |
| BP558 | CGGCATTCATTACCTCTGAGCTCAAG | DS4 delta fliQ detect, R, 250bp downstream |
| BP559 | TGGTCAATCAAACCAGCGTTCAAG | DS4 delta fliP F1 |
| BP560 | gctaattcagtttaagcggccatGATTGGCACAGAGAGAAGAGGGTAAG | DS4 delta fliP R1, universal overlap homology |
| BP561 | atggccgcttaaactgaattagcGGCGGATTGAGTTATCCTTTGTCATG | DS4 delta fliP F2, universal overlap homology |
| BP562 | CTCAGCAGCAAGCGCGTGTAGATG | DS4 delta fliP R2 |
| BP563 | TGTTGAAACGCATGCAAGTGCCAG | DS4 delta fliP detect, R, 250bp downstream |
| BP564 | CAGAACTTGTGCCACGGCAGTAAATAAAC | DS4 delta fliO F1 |
| BP565 | gctaattcagtttaagcggccatCATGACAAAGGATAACTCAATCCGCC | DS4 delta fliO R1, universal overlap homology |
| BP566 | atggccgcttaaactgaattagcGCTTCAATCGAGTTTTGATTGAACTGC | DS4 delta fliO F2, universal overlap homology |
| BP567 | CCGGTACAAGAATTAGCGCAATTCG | DS4 delta fliO R2 |
| BP568 | CGACACCATCATGGACATCCCAGTC | DS4 delta fliO detect, R, 250bp downstream |
| BP569 | ACGGAATATCGATCGCCACGACG | DS4 delta fliN F1 |
| BP570 | gctaattcagtttaagcggccatTGATCAGCCAAACTGAACGCATC | DS4 delta fliN R1, universal overlap homology |
| BP571 | atggccgcttaaactgaattagcTTTTCTGTCCTGTGTTAAACCTTACTTGC | DS4 delta fliN F2, universal overlap homology |
| BP572 | AAAAGAAACATTGGATGCAAATGGTAAACC | DS4 delta fliN R2 |

| | | |
|---|---|---|
| BP573 | ACTTACCGCGTGAAGATGGGGC | DS4 delta fliN detect, R, 250bp downstream |
| BP574 | ACGATCAGCACTGATGCCGATGC | DS4 delta fliM F1 |
| BP575 | gctaattcagtttaagcggccatAAACCCTTACTCAATACCATCGCCTC | DS4 delta fliM R1, universal overlap homology |
| BP576 | atggccgcttaaactgaattagcTGGCTTAATAGATCGGTCACGTTATACC | DS4 delta fliM F2, universal overlap homology |
| BP577 | TTCTAAGCGTTTCTCATCGTTATATTGCG | DS4 delta fliM R2 |
| BP578 | AATCAAAGCCCAGCTGATGGTGC | DS4 delta fliM detect, R, 250bp downstream |
| BP579 | TTCGTTGATAGAGGTTGCCGCTTGG | DS4 delta fliL F1 |
| BP580 | gctaattcagtttaagcggccatGGTATAACGTGACCGATCTATTAAGCCAAG | DS4 delta fliL R1, universal overlap homology |
| BP581 | atggccgcttaaactgaattagcTTCTGCCATTACTGTCTCTTATTGTTCTG | DS4 delta fliL F2, universal overlap homology |
| BP582 | CATGAAAGAGTCTGTGCCTTATGACATG | DS4 delta fliL R2 |
| BP583 | AAATGTTGTCTCAGCAAGGCATGC | DS4 delta fliL detect, R, 250bp downstream |
| BP584 | TTTTCAATCACTGGCTTACCCACCG | DS4 delta fliJ F1 |
| BP585 | gctaattcagtttaagcggccatAGCGTTTCTCATCGTTATATTGCGTTAG | DS4 delta fliJ R1, universal overlap homology |
| BP586 | atggccgcttaaactgaattagcACGCTATTACCCACCCAAGATGCTC | DS4 delta fliJ F2, universal overlap homology |
| BP587 | CTGGTGGCTGCTCTGGGTGAG | DS4 delta fliJ R2 |
| BP588 | TTATGCCACAAATCACCACAGAAGAG | DS4 delta fliJ detect, R, 250bp downstream |
| BP589 | TTTTCCTTTTGGTGCATCCGCACC | DS4 delta fliI F1 |
| BP590 | gctaattcagtttaagcggccatATAGCGTATGAATAATGCGATGGAATTCC | DS4 delta fliI R1, universal overlap homology |
| BP591 | atggccgcttaaactgaattagcTTGCATGGACTACTCGCCCC | DS4 delta fliI F2, universal overlap homology |
| BP592 | ATTACGGTAACCGATCAACACGGTC | DS4 delta fliI R2 |
| BP593 | CGCTGCCTATTTCTGGTCATCC | DS4 delta fliI detect, R, 250bp downstream |
| BP594 | TGCGCCACGTTGCTTGTTTGTG | DS4 delta fliH F1 |
| BP595 | gctaattcagtttaagcggccatGTCCATGCAAGCCTTGGCCG | DS4 delta fliH R1, universal overlap homology |
| BP596 | atggccgcttaaactgaattagcCCATGCTTAACCTACAATCTGTCTATGTTG | DS4 delta fliH F2, universal overlap homology |
| BP597 | AGTGTTGCCTTCGATGCCACTGTAC | DS4 delta fliH R2 |

| | | |
|---|---|---|
| BP598 | GCGACGATATCGAAGCAATGCCTCC | DS4 delta fliH detect, R, 250bp downstream |
| BP599 | GCAAAACTGAACAATGGTTAACGTGG | DS4 delta fliG F1 |
| BP600 | gctaattcagtttaagcggccatATCCTTTCTTGTTCTTCCTGTTTAGGC | DS4 delta fliG R1, universal overlap homology |
| BP601 | atggccgcttaaactgaattagcTCGTTAGCCATTTTCGTTCATCCAG | DS4 delta fliG F2, universal overlap homology |
| BP602 | GAAAGGTGCATTTACTGGCGCGG | DS4 delta fliG R2 |
| BP603 | TGCTCAACCCAGCAAGTGGC | DS4 delta fliG detect, R, 250bp downstream |
| BP604 | GCCATTTCACGAGACAAGACAATGTG | DS4 delta fliF F1 |
| BP605 | gctaattcagtttaagcggccatAATGGCTAACGATATCGTACCTCAAGATG | DS4 delta fliF R1, universal overlap homology |
| BP606 | atggccgcttaaactgaattagcGCCACACTTATTACCTAAAATTACACTGGC | DS4 delta fliF F2, universal overlap homology |
| BP607 | ATAGCTTGGTACGCTAATTGCTTGC | DS4 delta fliF R2 |
| BP608 | GTGCGAAAGTTGGGGCAGATTTTGG | DS4 delta fliF detect, R, 250bp downstream |
| BP609 | AATTGTTATCGACAAAAGCCGCTTACTG | DS4 delta fliS F1 |
| BP610 | gctaattcagtttaagcggccatGCAGCAGAAGTCGGCATTTAATCG | DS4 delta fliS R1, universal overlap homology |
| BP611 | atggccgcttaaactgaattagcCAAAGAACCGCGCATAGTATTCCTC | DS4 delta fliS F2, universal overlap homology |
| BP612 | TCGAAGAGCCGTATTAAAGATACTGACTAC | DS4 delta fliS R2 |
| BP613 | TTAATACTGAAGAAATCCTTCACTTGGTCG | DS4 delta fliS detect, R, 250bp downstream |
| BP614 | GCGCGGTTTCGACCATTGGC | DS4 delta fliT F1 |
| BP615 | gctaattcagtttaagcggccatAAAGAGGAATACTATGCGCGGTTCTTTG | DS4 delta fliT R1, universal overlap homology |
| BP616 | atggccgcttaaactgaattagcCTATTGGCCCAACGCGTTCATCAG | DS4 delta fliT F2, universal overlap homology |
| BP617 | ATCAACGGTCAAAGCGAAGATGTAAAAG | DS4 delta fliT R2 |
| BP618 | GCGGTAACACAGGTTTCGCGAAAC | DS4 delta fliT detect, R, 250bp downstream |
| BP619 | AACGCGTTTCCGTTAGATCGGTGATC | DS4 delta fliD F1 |
| BP620 | gctaattcagtttaagcggccatGGCCAATAGATGAAAGATGCACTCATAG | DS4 delta fliD R1, universal overlap homology |
| BP621 | atggccgcttaaactgaattagcGGGCCTAAACTCATCAAATCACCTC | DS4 delta fliD F2, universal overlap homology |
| BP622 | TTTGTCTTCAGGTTTCAAAATCAACAGC | DS4 delta fliD R2 |

| | | |
|---|---|---|
| BP623 | AAAATGGTCGAGCAAATGAATGAATTTGTG | DS4 delta fliD detect, R, 250bp downstream |
| BP624 | TTTGCGCAATTGAGTGTTAGAAAGGG | DS4 delta flaG F1 |
| BP625 | gctaattcagtttaagcggccatCGGTTTGTTAGTTGAAAAGGTGTAAGTCG | DS4 delta flaG R1, universal overlap homology |
| BP626 | atggccgcttaaactgaattagcTACAATCTCCCTTCCACCTTATGAGC | DS4 delta flaG F2, universal overlap homology |
| BP627 | AATGGTGAAGCAGCAGATCTTACAAC | DS4 delta flaG R2 |
| BP628 | CTAGACAACATTAACGAGAACGTGAACG | DS4 delta flaG detect, R, 250bp downstream |
| CAS0158 | GTGGCTGTTCGTAACGCGAAC | F1 primer delta FlaE (DSB67_RS03825) |
| CAS0202 | GTTCATCGCTGCAACCTATTCAGCTAATTCAGTTTAAGCGGCCAT | R1 for delta FlaE SOE (DSB67_RS03825) |
| CAS0160 | atggccgcttaaactgaattagcTGAATAGGTTGCAGCGATGAACATCG | F2 primer delta FlaE (DSB67_RS03825) |
| CAS0161 | CGTAACCGTTCTACCAAGCTCTGG | R2 primer delta FlaE (DSB67_RS03825) |
| CAS0162 | CGTAACTCTAACTTCGCAGAAGTG | R delta FlaE detect primer |
| CAS0163 | GGATGCCGAACATGGTGATCGTC | F1 primer delta FlaF (DSB67_RS11440) |
| CAS0164 | gctaattcagtttaagcggccatCAGCATATCGGTTGCACTGGTC | R1 primer delta FlaF (DSB67_RS11440) |
| CAS0165 | atggccgcttaaactgaattagcCTTGGTTAATGATTCGCACCACACG | F2 primer delta FlaF (DSB67_RS11440) |
| CAS0166 | CCAGAGAATTCAACGTCACCTTCAAC | R2 primer delta FlaF (DSB67_RS11440) |
| CAS0189 | CCAAGTAGTTCTCTCAAAGTTAC | R delta FlaF detect primer |
| CAS0168 | CGGTACTTACGGTACTCAGTC | F1 primer delta FlaA (DSB67_RS11425) |
| CAS0169 | gctaattcagtttaagcggccatCGCCATAGTTGATCTCCTTAAGGC | R1 primer delta FlaA (DSB67_RS11425) |
| CAS0170 | atggccgcttaaactgaattagcGGCTAATTGGGTCGACTCAGCC | F2 primer delta FlaA (DSB67_RS11425) |
| CAS0171 | GAAGCAATGTCACCACCATCTTCC | R2 primer delta FlaA (DSB67_RS11425) |
| CAS0172 | CGATTCTTCATCGACTCGAAACGC | R delta FlaA detect primer |
| CAS0192 | CGTGATTATGACGGTAGCGTGG | F1 delta FlaB (DSB67_03820) |
| CAS0193 | gctaattcagtttaagcggccatCACTGCCATGGTGATTTCTCCAATTG | R1 delta FlaB (DSB67_03820) |
| CAS0194 | atggccgcttaaactgaattagcCTAGGTTAATAAACCTAATCGACGTG | F2 delta FlaB (DSB67_03820) |
| CAS0195 | CCTATTCATGGACCGTGATGCG | R2 delta FlaB (DSB67_03820) |

| | | |
|---|---|---|
| CAS0196 | CACCAACATAATGCTGTCCTC | delta FlaB detect primer |
| CAS0197 | GGAGTGAGTAGAGACAGCGGTAG | F1 delta FlaD (DSB67_RS11430) |
| CAS0198 | gctaattcagtttaagcggccatCACTGCCATGGTGATTTCTCC | R1 delta FlaD (DSB67_RS11430) |
| CAS0199 | atggccgcttaaactgaattagcCCTAACTCAGCGCTAAGTCTTCTAGGT | F2 delta FlaD (DSB67_RS11430) |
| CAS0200 | TCACGTACGAATCGGTTAGCGTTC | R2 delta FlaD (DSB67_RS11430) |
| CAS0201 | CGCCATAGTTGATCTCCTTAAGGC | delta FlaD detect primer |
| **qPCR Primers** | | |
| BP345 | ATGGCTAAGGGGCAATCTCTAC | qRT-PCR DS40M4 Hfq F |
| BP346 | CTTGCAGTTTGATACCGTTCAC | qRT-PCR DS40M4 Hfq R |
| BP330 | ATGGCGATTAACGTTAATACTAACGTTTC | qRT-PCR DS40M4 flaA F |
| BP331 | AAGACAAACGCTCCATTGATTTTTGTTG | qRT-PCR DS40M4 flaA R |
| BP358 | ATGAATGTGAATTTATCCAATGTTTCTG | qRT-PCR DS40M4 fliK F |
| BP359 | AAAAACCTTTGCTTTCAGGGT | qRT-PCR DS40M4 fliK R |
| BP650 | ATGAAAAGATGGCTTGTTGCCG | qRT-PCR DS40M4 flgO F |
| BP651 | ATTGGCTGCCTGAATACGGTTC | qRT-PCR DS40M4 flgO R |
| BP673 | AGCACAACCAAAATACATGATTGCG | qRT-PCR DS40M4 flhF F |
| BP674 | CTAGAGTCCTTCGCTGTCACTG | qRT-PCR DS40M4 flhF R |
| BP675 | TTTAGACTTTGCTAAGGAATTGCAGG | qRT-PCR DS40M4 flgB F |
| BP676 | GACCAAGGTTGTAGTGGCAGG | qRT-PCR DS40M4 flgB R |

### References

1. Bassler BL, Greenberg EP, Stevens AM. 1997. Cross-species induction of luminescence in the quorum-sensing bacterium Vibrio harveyi. J Bacteriol 179:4043 LP – 4045.
2. Dias GM, Thompson CC, Fishman B, Naka H, Haygood MG, Crosa JH, Thompson FL. 2012. Genome sequence of the marine bacterium Vibrio campbellii DS40M4, isolated from open ocean water. J Bacteriol 194:904.
3. Simpson CA, Podicheti R, Rusch DB, Dalia AB, van Kessel JC. 2019. Diversity in Natural Transformation Frequencies and Regulation across Vibrio Species. MBio 10:1–16.
4. Tschirhart T, Shukla V, Kelly EE, Schultzhaus Z, NewRingeisen E, Erickson JS, Wang Z, Garcia W, Curl E, Egbert RG, Yeung E, Vora GJ. 2019. Synthetic Biology Tools for the Fast-Growing Marine Bacterium Vibrio natriegens. ACS Synth Biol 8:2069–2079.
